# StocSum: stochastic summary statistics for whole genome sequencing studies

**DOI:** 10.1101/2023.04.06.535886

**Authors:** Nannan Wang, Bing Yu, Goo Jun, Qibin Qi, Ramon A. Durazo-Arvizu, Sara Lindstrom, Alanna C. Morrison, Robert C. Kaplan, Eric Boerwinkle, Han Chen

**Author notes:** Correspondence: Han Chen.

## Abstract

Genomic summary statistics, usually defined as single-variant test results from genome-wide association studies, have been widely used to advance the genetics field in a wide range of applications. Applications that involve multiple genetic variants also require their correlations or linkage disequilibrium (LD) information, often obtained from an external reference panel. In practice, it is usually difficult to find suitable external reference panels that represent the LD structure for underrepresented and admixed populations, or rare genetic variants from whole genome sequencing (WGS) studies, limiting the scope of applications for genomic summary statistics. Here we introduce StocSum, a novel reference-panel-free statistical framework for generating, managing, and analyzing stochastic summary statistics using random vectors. We develop various downstream applications using StocSum including single-variant tests, conditional association tests, gene-environment interaction tests, variant set tests, as well as meta-analysis and LD score regression tools. We demonstrate the accuracy and computational efficiency of StocSum using two cohorts from the Trans-Omics for Precision Medicine Program. StocSum will facilitate sharing and utilization of genomic summary statistics from WGS studies, especially for underrepresented and admixed populations.

## Main

International consortia for genomic epidemiology research on complex diseases and quantitative traits have generated a great abundance of genomic summary statistics^1–11^. These summary statistics are often in the form of regression coefficients and their standard errors (and/or z scores) from single-variant tests for common genetic variants, typically defined as those with a minor allele frequency (MAF) of greater than 5% or 1%, in genome-wide association studies (GWAS). Genomic summary statistics contain important information for researchers without direct access to individual-level genotype data and sharing genomic summary results is now commonly mandated by scientific journals and funding agencies. Genomic summary statistics also play a crucial role for cross-institutional (both national and international) collaborations where individual-level data are difficult to share due to ethical and legal restrictions.

Genomic summary statistics have been used to address different scientific questions in genetic and genomic research, such as meta-analysis^12,13^, heritability estimation^14–16^, conditional analysis^17^, variant set^18–21^ and gene-based tests^22,23^, multiple phenotype analysis^24–26^, genetic correlation or co-heritability estimation^27,28^, and others^29,30^. Many of these methods also require information on the linkage disequilibrium (LD) or correlation structure between genetic variants, which is commonly derived from external reference panels^14–17,23^. While these methods usually have good performance for common variants in populations of European ancestry, it has been challenging to extend the scope of summary statistic-based applications to other ancestry groups and admixed populations^14^ as well as rare variants^15^, defined as those with MAF < 5% or 1%, since the LD patterns in an external reference panel often do not match with those in the study sample.

Current large-scale whole genome sequencing (WGS) projects, such as the National Heart, Lung, and Blood Institute’s (NHLBI’s) Trans-Omics for Precision Medicine (TOPMed) program, the National Human Genome Research Institute’s (NHGRI’s) Centers for Common Disease Genomics (CCDG) initiative, and the National Institute on Aging’s (NIA’s) Alzheimer’s Disease Sequencing Project (ADSP), have unveiled hundreds of millions of rare variants from diverse populations. Making efficient and flexible use of these WGS resources and derived genomic summary results is paramount to facilitate international collaborations and scientific discoveries. However, managing and coordinating large-scale consortium efforts on rare variant meta-analyses has been quite challenging, since many existing meta-analysis software programs such as seqMeta^31^, MetaSKAT^18^, RVTESTS^32^, RAREMETAL^33^ and SMMAT^21^, require the correlation (or LD) matrices for rare variants to be computed internally in the study samples. In rare variant tests^21,34–40^, variant set definitions often need to be pre-specified (e.g., by genomic motifs such as genes or physical windows). Therefore, researchers have to recreate the LD matrices every time they want to redefine a variant set (e.g., by including more variants in a test region or combining two testing windows). This requires additional computational resources, making it difficult for researchers to efficiently leverage the richness of the data. On the other hand, sharing terabytes or even petabytes of individual-level WGS and phenotype data across research groups is a daunting task, and the risk of privacy breaches generally increases as more copies of individual-level data are being shared. Although individual-level WGS data can now be accessed through cloud-based computing platforms such as the Analysis Commons^41^, BioData Catalyst and AnVIL, and recently developed analysis tools such as STAARpipeline^42^ have greatly improved rare variant analyses especially for the noncoding genome, research groups are still largely constrained by the computational costs they can afford in running WGS data analysis using individual-level data directly.

Ideally, computing genomic summary statistics only once and then recycling them for different variant set definitions and weighting schemes is a more efficient strategy for WGS analysis on rare variants. Downstream analyses using summary statistics would not depend on the sample size *N* and therefore could be easily performed on a desktop computer. However, there are critical barriers in scaling existing statistical methods based on GWAS summary statistics up to allow for summary statistics based on WGS studies. First, calculating traditional pairwise LD measures from individual-level genomic data is computationally intensive. In general, a covariance matrix of size *M* × *M* is desired (**Fig.1a**), where *M* is the total number of variants, which has already exceeded 700 million in TOPMed. In practice, genotype data are usually saved by chromosome, but *M* is still on the scale of millions even for the shortest chromosome, making pairwise LD calculations on the whole genome (or one chromosome) computationally infeasible. Second, although restricting LD calculations to only genetic variants in close proximity (e.g., the sliding window strategy^43^ and the banded sparse LD matrices in 500kb windows^44^) is more computationally efficient than calculating the full *M* × *M* covariance matrix, it does not allow for the flexibility of testing distant genetic variants jointly. As there is growing evidence that the three-dimensional organization of chromosomes profoundly affects gene regulation^29,45–52^, LD matrices generated through sliding windows cannot be used if the variant set of interest contains genetic variants that are located far away from each other. In addition, LD statistics used in rare variant tests can greatly depend on the phenotype of interest (e.g., the phenotype distributions in minor allele carriers vs. non-carriers for each variant), and generally cannot be pre-computed using WGS data without the phenotype information.

**Figure 1:**
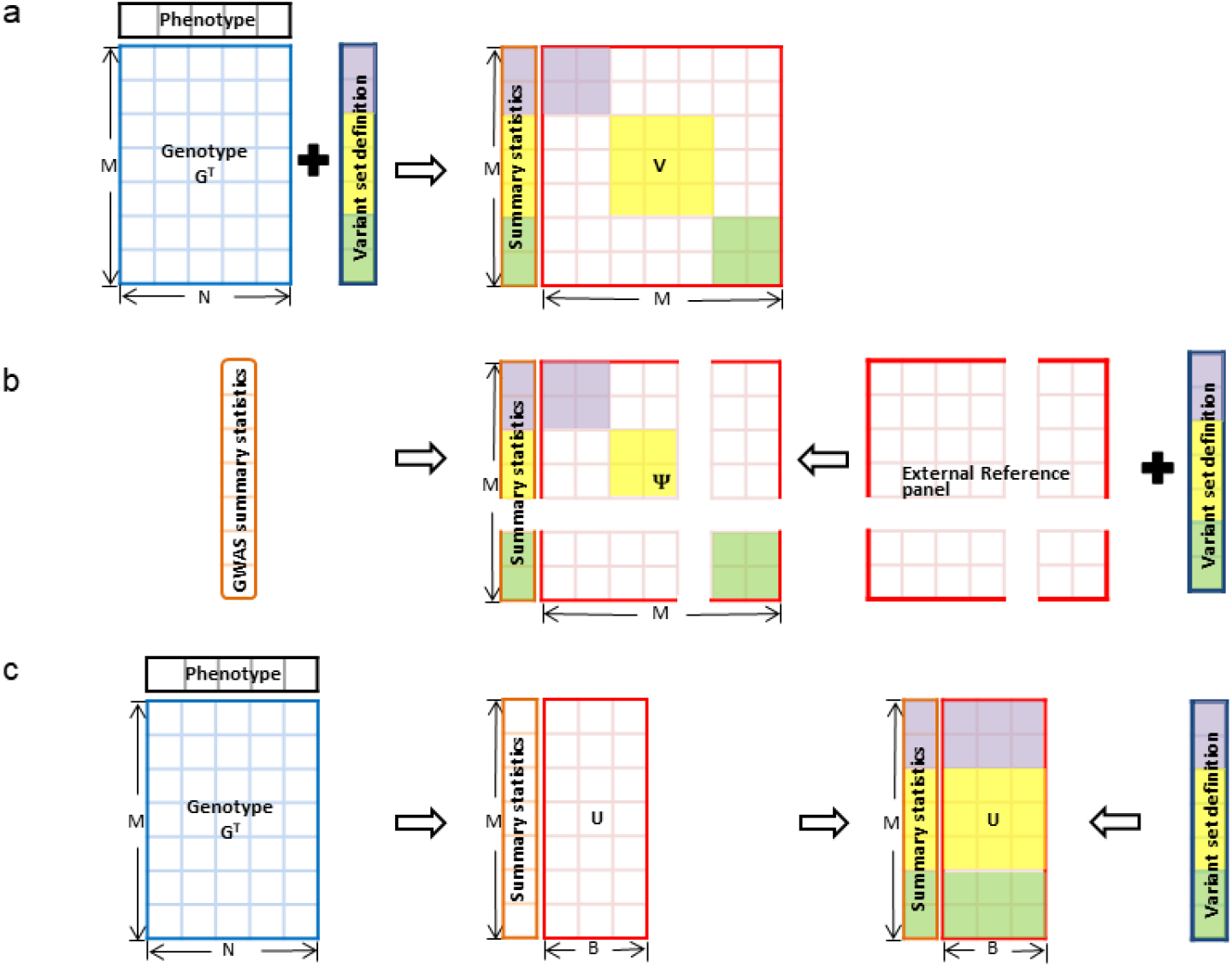
The StocSum framework. **a)** Traditional methods calculate the correlation or LD matrix ***V*** from individual-level genotype data. To reduce the computational burden, the full *M* × *M* matrix is usually not computed in practice, but rather replaced by a block-diagonal or banded sparse matrix based on pre-defined variant sets, at the cost of losing the flexibility in testing distant genetic variants jointly. **b**) The approximate LD matrix ***ψ*** is obtained from external reference panels when individual-level genotypes are not available, in many genomic summary statistic-based applications. However, variants may be excluded if they do not exist in the reference panel. **c**) StocSum generates stochastic summary statistics ***U*** from random vectors, which can be used to efficiently look up the covariance among arbitrary variant sets that are not pre-defined. *M*, the number of variants. *N*, the sample size. *B*, the number of random vectors used to construct stochastic summary statistics ***U***.

In addition, many existing methods using genomic summary statistics based on common variants rely on LD information from external reference panels^14–17,23^ (**Fig. 1b**). These methods have been widely applied to common variants in primary populations of European ancestry. Extension of these methods to underrepresented and admixed populations, however, has been noted as a challenge^14,27^ due to lack of appropriate reference panels that accurately represent the LD structure.

In this study, we propose the StocSum framework as illustrated in **Fig. 1c** to extend the scope of summary statistic-based applications. For methods that require between-variant correlation or LD matrices, we use a stochastic summary statistic matrix ***U*** to replace the traditional pairwise LD matrix ***V***. Specifically, by using *B* independent and identically distributed random vectors to represent the parametric distribution of any model-based residuals from a complex statistical model that accounts for potential sample correlations, matrix ***U*** can be quickly computed by matrix multiplication of the *N* × *M* genotype matrix ***G*** and these *B* random vectors. The size of ***U*** scales linearly with *M* and *B* (i.e., *O*(*MB*)), compared to quadratically in the form of a traditional pairwise LD matrix ***V***. The stochastic summary statistic matrix ***U*** can always be computed in linear time with the sample size *N* (*i.e., O*(*NMB*)), regardless of any complex sample correlation structures, compared to *O*(*NM*^2^) for the traditional pairwise LD matrix ***V*** in classical linear and logistic regression models for unrelated individuals, or mixed effect models to account for sample correlations in the presence of a sparse and block-diagonal relatedness matrix with bounded block sizes (e.g., a population-based family study, with known pedigrees). The complexity for computing ***V*** could further increase to *O*(*N*^2^*M* + *NM*^2^) if the relatedness matrix used in the mixed effect model is not block-diagonal (e.g., the genetic relationship matrix, or GRM). We also develop downstream applications using StocSum, including single-variant, conditional association, gene-environment interaction, variant set tests, as well as meta-analysis and LD score regression tools. This framework can flexibly accommodate changes of variant set definitions in analysis plans. For example, in variant set tests for rare variants, we can efficiently calculate the LD matrix for any variant sets by simply looking up 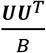 rather than rerunning the analysis with individual-level genotype data to update LD matrices for new variant sets. Compared with using external reference panels which might not well represent the LD structure in study samples from underrepresented and admixed populations, StocSum can be used to better calibrate the LD information in a wide range of genomic summary statistic-based applications.

## Results

### Overview of the method

We describe StocSum under the generalized linear mixed model (GLMM) framework. It can also be applied to simpler statistical models such as generalized linear models^53^ and extended to more complex models such as generalized additive mixed models^54^. The GLMM can be written as:

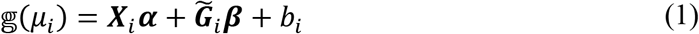

where 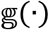 is a monotonic link function of *μ_i_*, and 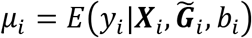 is the conditional mean of the phenotype *y_i_* given *p* covariates ***X***_*i*_, *q* genotypes 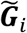 and random effects *b_i_*, for individual *i* of *N* samples. The phenotype *y_i_* follows a distribution in the exponential family, such as a normal distribution for continuous phenotypes, or a Bernoulli distribution for binary phenotypes. Here ***α*** is a length *p* column vector of fixed covariate effects including an intercept term. The genotype matrix 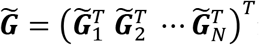 is an *N* × *q* matrix for *q* (*q* ≥ 1) genetic variants and ***β*** is a length *q* genotype effect vector. We assume that ***b*** = (*b*_1_ *b*_2_ ⋯ *b_N_*)^*T*^ is a length *N* column vector of random effects and 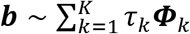, where *τ_k_* are the variance component parameters and ***Φ***_*k*_ are known *N* × *N* dense or sparse relatedness matrices which account for multiple layers of correlation structures, such as genetic relatedness, hierarchical designs, shared environmental effects and repeated measures from longitudinal studies.

For both single-variant (*q* = 1) and variant set (*q* > 1) tests, we only need to fit the null model 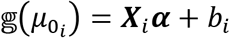 without fixed genetic effects one time, then each test can be constructed using single-variant scores ***S*** and *q* × *q* covariance matrices 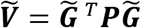, where ***P*** is the projection matrix from this model^21,55^. Denote *M* as the total number of genetic variants on the whole genome (or one chromosome). To avoid computing the full *M* × *M* matrix ***V*** or its block-diagonal version for every *q* variants 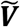 directly from individual-level data or an external reference panel, StocSum leverages a length *N* random vector ***R***_*b*_ from a multivariate normal distribution with mean **0** and covariance matrix ***P***.

Then it repeats this simulation process *B* times and combines these random vectors into an *N* × *B* random matrix ***R*** = (***R***_1_ ***R***_2_ ⋯ ***R***_*B*_). Denoting ***U*** = ***G***^*T*^***R*** as the stochastic summary statistics for *M* genetic variants on the whole genome (or one chromosome), for arbitrary *q* variants (*q* < *B*), we can extract the corresponding rows from the *M* × *B* stochastic summary statistics matrix ***U*** as 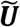 and use 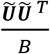 to estimate the covariance matrix 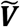.

To implement StocSum and various downstream genetic analysis applications, our framework comprises four major steps: (1) fitting a generalized linear mixed model under the null hypothesis, e.g., 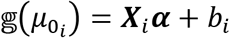, estimating variance component parameters, residuals and the projection matrix ***P***; (2) generating an *N* × *B* random matrix ***R***, with each column of ***R*** simulated from a multivariate normal distribution with mean **0** and covariance matrix ***P***; (3) using individual-level genotypes ***G*** to compute score statistics from residuals, and the stochastic summary statistics matrix ***U*** = ***G***^*T*^***R***; and (4) computing *P* values in each downstream application (see **Methods**). The first three steps could be shared by multiple genetic analysis applications including single-variant, conditional association, gene-environment interaction, and variant set tests. We could also estimate LD scores efficiently in the stochastic summary statistics framework, thus extending its application to underrepresented and admixed populations (see **Methods**).

### Single-variant tests

To evaluate the performance of StocSum in single-variant tests, we used TOPMed WGS freeze 8 data from the Hispanic Community Health Study/Study of Latinos (HCHS/SOL). After quality control we had data for 120M variants in 7,297 individuals (**Methods**). We first compared *P* values calculated by StocSum with different numbers of random vector replicates *B* and GMMAT^55^ using individual-level genotypes in a genome-wide single-variant analysis of blood low-density lipoprotein (LDL) cholesterol levels (**Fig. 2a-d**). The *P* values calculated from StocSum were compared with those from GMMAT using individual-level data. No systematic genomic inflation was observed from the quantile-quantile (Q-Q) plots (**Fig. S1**). StocSum *P* values were close to GMMAT when the number of random vector replicates *B* ranged from 100 to 10,000 (Fig.**2b-2d**). We did observe that a small *B* (*B*=10) led to inaccurate *P* values (**Fig. 2a**).

**Figure 2:**
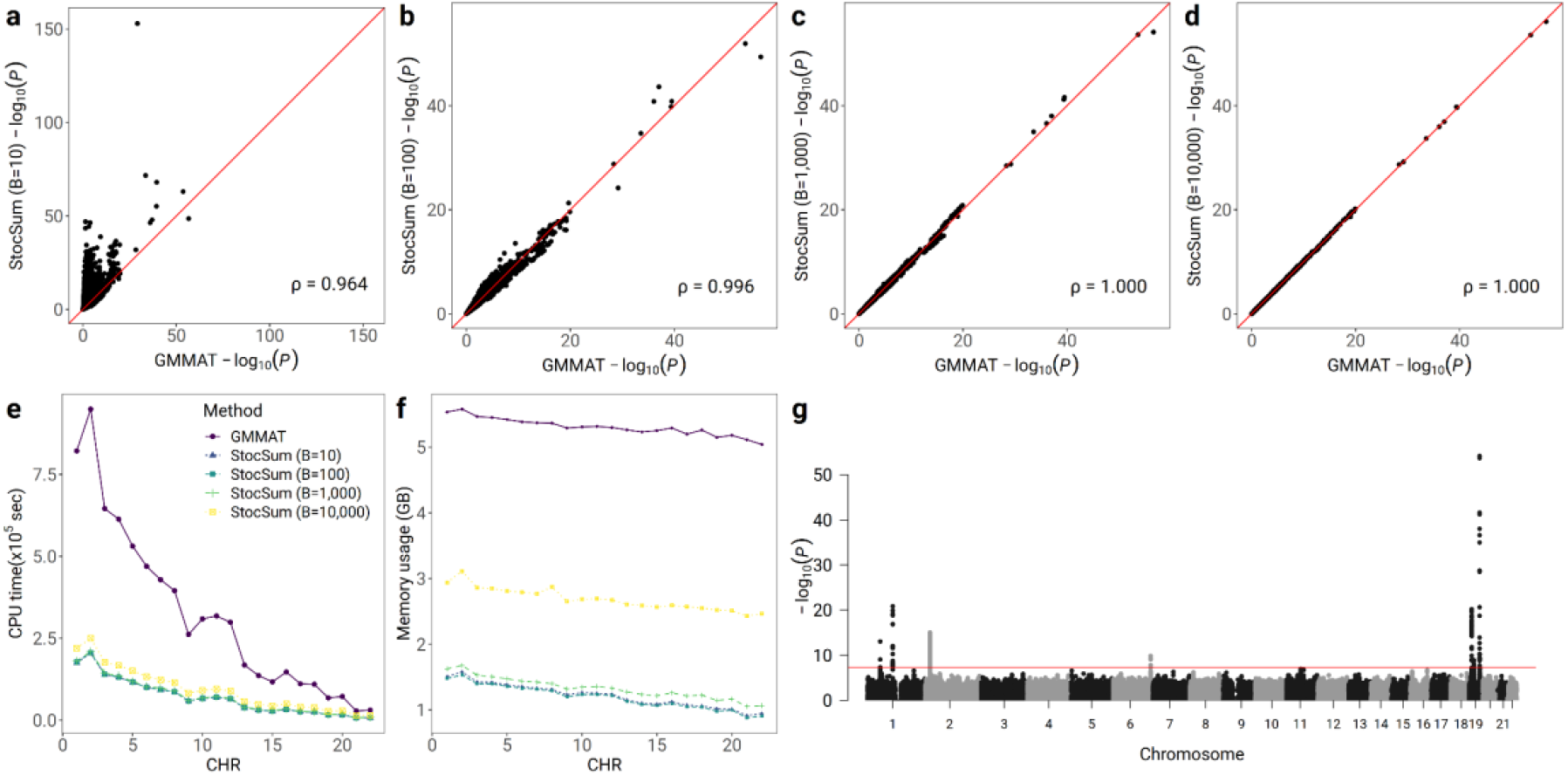
StocSum in single-variant tests. **a-d**) comparison of *P* values from GMMAT and StocSum with the number of random vector replicates *B* being equal to 10 **(a)**, 100 **(b)**, 1,000 **(c)** and 10,000 **(d)**. The x axis and the y axis represent −log_10_(*P*) from single-variant tests using GMMAT and StocSum, respectively. The red line denotes the reference line of equality. Spearman’s rank correlation coefficients are shown at the bottom right. **e**) comparison of CPU time between GMMAT and StocSum. The x axis represents the chromosome numbers, and the y axis represents the CPU time in 10^5^ seconds. For GMMAT, the CPU time consists of fitting the null model and conducting the association test. For StocSum, the CPU time is the sum of four steps: fitting the null model, generating the random vectors, computing the single-variant score statistics and the stochastic summary statistics, and computing the *P* values. **f**) comparison of memory usage by GMMAT and StocSum. The x axis represents the chromosome numbers and the y axis represents the memory footprint per core in GB. The data used in this test consisted of 120M variants from 7,297 individuals in HCHS/SOL. All tests were performed on a high-performance computing server, with 64 cores used for parallel computing. **g**) the Manhattan plot of single-variant test on LDL in the HCHS/SOL study using StocSum. The x-axis represents the physical chromosome and position of each variant and the y-axis represents −log_10_(*P*) from the StocSum single-variant test. Only variants with MAF > 0.5% were included in the Manhattan plot. The red line indicates the genome-wide significance level on the log scale, −log_10_ (5×10^−8^).

To demonstrate the computational efficiency of StocSum, we ran GMMAT and StocSum (*B*=1,000) on the same computing platform where 64 cores were used in parallel computing for both programs. Both runtime and memory usage of StocSum were much lower compared to GMMAT. For example, it took about 50.2 CPU hours to run chromosome 1 with 9.7M variants using StocSum, which was 4.6-fold faster than GMMAT. Meanwhile, StocSum only had 29.3% of the memory footprint compared to GMMAT. Across all 22 autosomes, StocSum was 4.4-fold faster than GMMAT, with about 25.1% of the memory footprint compared to GMMAT (**Fig. 2e-f**). As expected, both the run time and memory footprint increased with a larger *B*. However, the run time and memory footprint of StocSum when *B*=10,000 were still only 29.3% and 50.6% compared to GMMAT, respectively.

Using StocSum, we ran a WGS study of LDL cholesterol in HCHS/SOL and identified seven genome-wide significant (*P* values < 5×10^−8^) regions mapped to genes *PCSK9* and *CELSR2* on chromosome 1, *APOB* on chromosome 2, *LPA* on chromosome 6, *LDLR*, *SUGP1*, and *APOE* on chromosome 19 (**Fig. 2g**, **Table S1**), all of which had been previously reported to be associated with LDL^4,56–59^.

We also compared StocSum with fastGWA^60^, another widely used single-variant test tool (**Figs. S2–3**). To make a fair comparison on the same statistical model, we only included one random effect term for genetic relatedness, without allowing for heteroscedasticity in the null model for GMMAT and StocSum. Both fastGWA and GMMAT results were very similar (**Figs.S2–3**). In this different null model, StocSum *P* values were still consistent with GMMAT when *B* ranged from 100 to 10,000. The CPU time used by fastGWA was generally stable for different chromosomes (**Fig.S4a**). The total CPU time for the whole genome analysis was similar for StocSum (*B*=1,000) and fastGWA. The memory usage of fastGWA was slightly larger (about 1.7-fold) compared to StocSum with *B*=1,000 (**Fig.S4b**).

### Conditional association tests

We implemented StocSum for conditional association tests and applied it to the seven genome-wide significant regions identified in **Fig. 2g**. The sentinel variant in the *APOE* gene region is chr19: 44908822 (rs7412) with *P* = 7.1×10^−55^. There are 26 common variants with MAF > 0.5% close to this sentinel variant in this region, with a *P* value less than 5×10^−8^ (**Fig. 3**). After conditioning on the sentinel variant, we identified a secondary association variant chr19: 44908684 (rs429358) with conditional *P* = 8.2×10^−15^. After conditioning on both rs7412 and rs429358, all other variants in the region had *P* values > 1.8×10^−3^, indicating that no additional independent associations exist in this region. We also observed similar patterns in the other six regions (**Table S1**), after conditioning on either one or two top associated variants in each region (**Fig. S5 a-f**).

**Figure 3:**
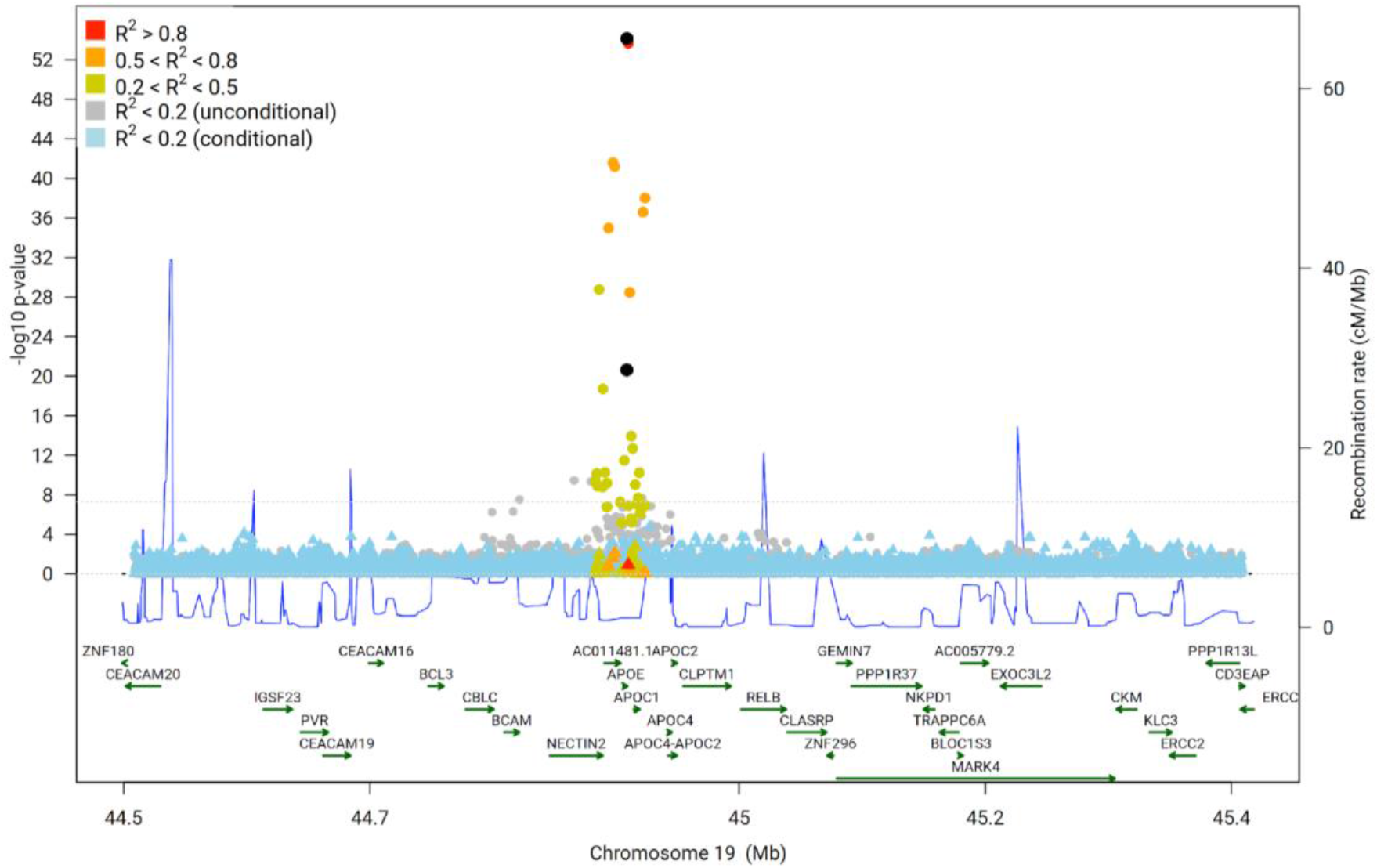
A regional plot of StocSum conditional association test results in the *APOE* region. Variants with MAF > 0.5% in a 1Mb window near association variants rs7412 and rs429358 (highlighted in black dots). Original single variant test *P* values are shown in dots and conditional *P* values are shown in triangles. Variants in four LD categories are shown in different colors based on the maximum squared correlation to the sentinel variant rs7412 and the secondary association variant rs429358 calculated in HCHS/SOL. The horizontal dashed line indicates the genome-wide significance level on the log scale, −log_10_ (5 × 10^−8^). The blue curve shows recombination rates from all populations in the 1000 Genome Project.

### Gene-environment interaction tests

We next developed and implemented a one-degree-of-freedom gene-environment interaction test and a two-degree-of-freedom joint test of the genetic main effects and the gene-environment interactions in the StocSum framework. We benchmarked our tests with MAGEE using individual-level data^61^. No systematic genomic inflation was observed from the quantile-quantile (Q-Q) plots (**Fig. S6**). **Fig. S7** shows *P* values from a gene-sex interaction analysis on waist-hip ratio (WHR) in HCHS/SOL. MAGEE and StocSum *P* values were highly consistent, with Spearman’s correlations of 1.000, 0.998, 0.999, respectively, for the marginal genetic effect test, the gene-environment interaction test and the joint test. We identified four potential loci from marginal genetic effect tests, three with significant gene-sex interactions, and four from the joint tests, at the suggestive significance level of 5 × 10^−7^, including six previously reported genome-wide significant loci in gene regions *COBLL1*, *IGF2R*, *AOAH*, *IQSEC3*, *TEKT5*, and *MAPT*^62–68^ (Table S2).

### Variant set tests

We also used TOPMed WGS freeze 8 data and LDL cholesterol levels from the HCHS/SOL study to illustrate variant set tests in the StocSum framework. We compared *P* values calculated by StocSum with different numbers of random vector replicates *B* and SMMAT^21^ using individual-level genotypes in a genome-wide 20 kb non-overlapping sliding window analysis on all genetic variants, using a beta density weight on the MAF with parameters 1 and 25. We noted that 20 kb was probably wider than what was commonly used in WGS sliding window analyses^43^, but we chose this window size to evaluate the performance of StocSum variant set tests in an extreme scenario not in favor of StocSum, because there could be many windows with the number of variants *q* > *B*. In this case, 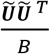 from StocSum would not be an appropriate estimate for the *q* × *q* covariance matrix 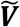 computed directly from individual-level data, since only *B* singular values could be computed from the *q* × *B* matrix 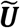.

**Figs. 4a-d** shows comparisons of *P* values from SMMAT using individual-level genotypes and StocSum with *B* ranging from 10 to 10,000. When *B*=1,000 or 10,000, *P* values from the two methods were highly consistent (**Figs. 4c-d**). For windows with small SMMAT *P* values, StocSum tended to overestimate these *P* values when *B*=10 or 100 (**Fig. 4a-b**), possibly because only 10 or 100 singular values from 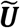 was insufficient to approximate the eigenvalues from the *q* × *q* covariance matrix 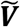 from SMMAT.

**Figure 4:**
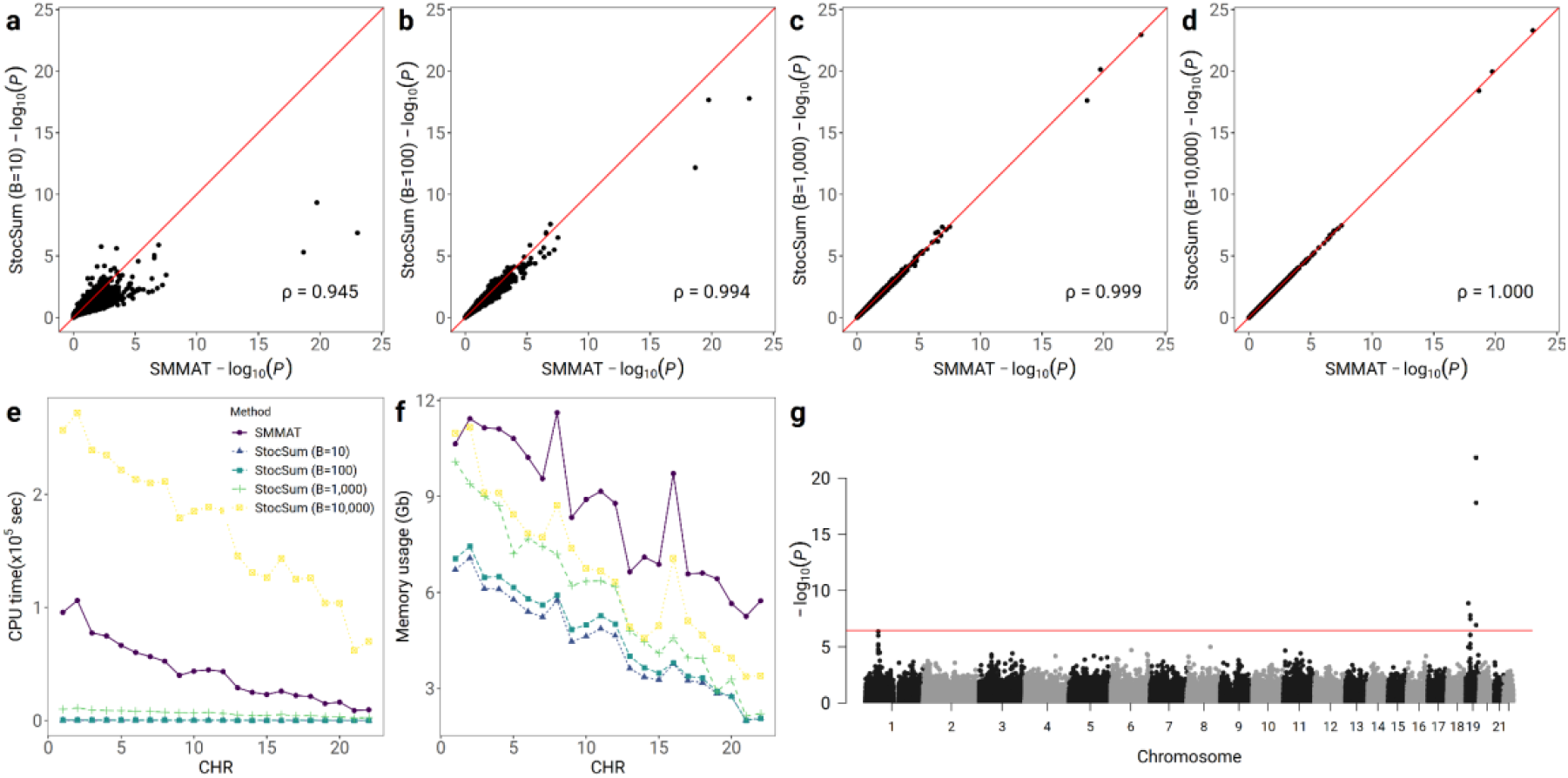
StocSum in variant set tests. Comparison of *P* values from SMMAT and StocSum with the number of random vector replicates *B* being equal to 10 (a), 100 (b), 1,000 (c) and 10,000 (d) in a 20 kb sliding window analysis on the whole genome. The x axis and the y axis represent −log_10_(*P*) from a whole genome 20 kb sliding window analysis, using variant set tests from SMMAT and StocSum, respectively, with a beta density weight on the MAF with parameters 1 and 25. The red line denotes the reference line of equality. Spearman’s rank correlation coefficients are shown at the bottom right. e, comparison of CPU time between SMMAT and StocSum. The x axis represents the chromosome numbers and the y axis represents the CPU time in 10^5^ seconds. For SMMAT, the CPU time did not include fitting the null model or reading the variant set definitions. For StocSum, the CPU time did not include computing stochastic summary statistics from individual-level data. f, comparison of memory usage by SMMAT and StocSum. The x axis represents the chromosome number and the y axis represents the memory footprint in GB. The data used in this test consisted of 120M variants from 7,297 individuals in HCHS/SOL, including all variants regardless of their MAF (such as singletons and doubletons). All tests were performed on a high-performance computing server, with a single thread for each chromosome. g, the Manhattan plot of 20 kb sliding window variant set tests on LDL in the HCHS/SOL study using StocSum. The x-axis represents the start physical chromosome and position of each variant set and the y-axis represents −log_10_(*P*) from the StocSum variant set test corresponding to SMMAT. The red line indicates the genome-wide significance level on the log scale, −log_10_ (3.7×10^−7^).

StocSum variant set tests are computationally efficient (**Figs. 4e-f**). It only took StocSum (*B*=1,000) 2.7 CPU hours to finish variant set tests on chromosome 1 using 20 kb sliding windows, which was 9.7-fold faster than SMMAT using individual-level data. Across the autosomes, there were a total of 134,739 non-overlapping 20 kb windows containing at least one variant. On average, the StocSum (*B*=1,000) CPU time was about 14.3% of the SMMAT CPU time. Meanwhile, StocSum (*B*=1,000) only required about 68.1% of the memory compared to SMMAT. StocSum with *B*=10,000 utilized more CPU time than SMMAT since *B* was larger than the sample size (N=7,297), making the *M* × *B* stochastic summary statistics matrix ***U*** even larger in size compared to the *N* × *M* genotype matrix ***G***. In this 20 kb sliding window analysis using StocSum variant set tests, we identified four regions associated with LDL levels in HCHS/SOL^4,56–59^, at the significance level of 0.05/134,739=3.7×10^−7^ (**Fig. 4g, Table S3**).

We next compared StocSum with fastBAT for variant set tests. fastBAT utilizes single-variant summary statistics from fastGWA and LD information from a reference panel such as the 1000 Genomes Project^69^. To make a fair comparison on the same statistical model and same weights used in variant set tests, we only included one random effect term for genetic relatedness, without allowing for heteroscedasticity in the null model for SMMAT and StocSum, and a beta density weight on the MAF with parameters 0.5 and 0.5 (which is equivalent to rescaling each variant with a unit variance as implemented in fastBAT). For fastBAT, we compared five different reference panels, including an internal reference panel using individual-level genotypes from the original study sample (called fastBAT (Sample)), as well as four external reference panels from the 1000 Genomes Project^69^: European populations (fastBAT (Eu)), European and African populations (fastBAT (EuAf)), European and American populations (fastBAT (EuAm)), and European, African and American populations (fastBAT (EuAfAm)). Variant set test *P* values from SMMAT, StocSum (*B*=1,000), and fastBAT (Sample) were highly concordant (**Figs. S9–10**), with pairwise Spearman correlation coefficients being greater than 0.99. However, fastBAT with external reference panels, i.e., fastBAT (Eu), fastBAT (EuAf), fastBAT (EuAm), fastBAT (EuAfAm), gave inaccurate variant set test *P* values compared to SMMAT using individual-level genotypes. The correlation coefficients of log_10_(*P*) between SMMAT and fastBAT with Eu, EuAf, EuAm, EuAfAm reference panels were 0.59, 0.77, 0.66, and 0.78, respectively (**Fig. S10**). Since Hispanic/Latino adults are three-way admixed populations with European, African and Amerindian ancestries, it is not surprising that an external reference panel from only European populations could not represent the LD structure in HCHS/SOL samples accurately. Interestingly, although including African and American populations in the external reference panel did improve the concordance of fastBAT *P* values compared to SMMAT, fastBAT using the internal reference panel clearly outperformed all external reference panels that we investigated. In addition, when an external reference panel was used, variants not included in the panel would have to be excluded, leading to loss of unique variants in the study samples. This highlights the importance of choosing an accurate reference panel for fastBAT, and the best reference panel for study samples from underrepresented, admixed or isolated populations are the study samples themselves. StocSum represents the LD structure in any variant sets through a stochastic summary statistic matrix ***U*** directly derived from study samples rather than external reference panels, thus providing accurate variant set test results. Meanwhile, StocSum with *B*=1,000 was slightly faster (1.7-fold) than fastBAT (Sample) on the whole genome (**Fig. S11a**), with a dramatically reduced memory footprint (3.6%) compared to fastBAT (Sample) (**Fig. S11b**).

To illustrate StocSum variant set tests beyond sliding windows, we compared StocSum (B=1,000) with SMMAT when the variant sets composed of different regions that were physically farther away. These variant sets were defined by merging chromatin loops of H3K27ac HiChIP interaction in the GM12878 cell line^70–72^. As the definition of variant sets changed, SMMAT required rerunning the analysis using individual-level genotypes, while StocSum variant set tests could directly extract information about these new variant sets from the same pre-computed stochastic summary statistic matrix ***U***, which yielded highly accurate *P* values (**Fig. S12a**), while using much less CPU time and memory (**Figs. S12b-c**).

### Meta-analysis

Meta-analysis in the StocSum framework can be performed by combining the stochastic summary statistic matrices ***U*** from different studies. To illustrate how single-variant and variant set tests can be conducted in a meta-analysis, we combined the stochastic summary statistic matrices ***U*** from three studies: longitudinal LDL levels as repeated measures in African-Americans (AA) from the Atherosclerosis Risk in Communities (ARIC) study (70M variants from 2,045 individuals) visits 1-6, European-Americans (EA) from ARIC (92M variants from 6,327 individuals) visits 1-6, and baseline LDL levels as cross-sectional measures in Hispanic/Latino adults from HCHS/SOL (120M variants from 7,297 individuals). *P* values from StocSum (*B*=1,000) were highly concordant with GMMAT results from longitudinal LDL level analyses, for both ARIC AA and EA subgroups (**Fig. S13**), which further demonstrated the robustness of StocSum in different populations. *P* values from StocSum meta-analysis (*B*=1,000) were highly concordant with those from GMMAT single-variant meta-analysis (**Fig. 5a**) and SMMAT variant set meta-analysis (**Fig. 5c**). We identified 14 LDL loci from StocSum meta-analysis (*B*=1,000) single-variant tests^4,56–59,73–76^ (**Fig. 5b, Table S4**), at the significance level of 5×10^−8^. In variant set tests (**Fig. 5d, Table S5**), we identified four regions associated with LDL levels from StocSum meta-analysis (*B*=1,000), at the significance level of 3.7×10^−7^.

**Figure 5:**
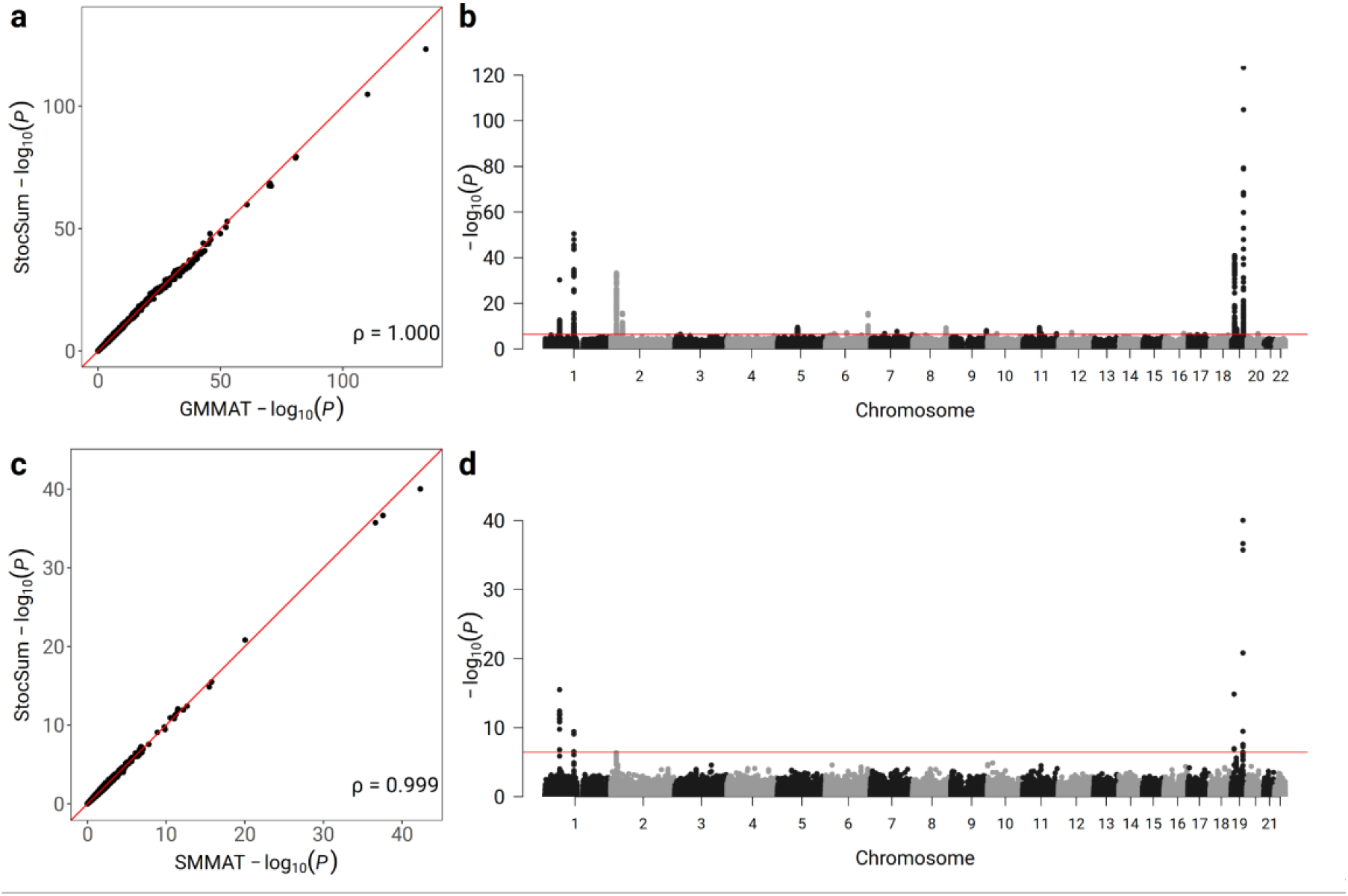
StocSum in meta-analysis. a, comparison of single-variant meta-analysis *P* values from GMMAT and StocSum with the number of random vector replicates *B* being equal to 1,000. The x axis and the y axis represent −log_10_(*P*) from single-variant meta-analysis using GMMAT and StocSum, respectively. The red line denotes the reference line of equality. Spearman’s rank correlation coefficients are shown at the bottom right. b, the Manhattan plot of single-variant tests on LDL in the meta-analysis of ARIC AA and EA, and HCHS/SOL studies using StocSum. The x-axis represents the physical chromosome and position of each variant and the y-axis represents −log_10_(*P*) from the StocSum single-variant test. Only variants with MAF > 0.5% were included in the Manhattan plot. The red line indicates the genome-wide significance level on the log scale, −log_10_ (5×10^−8^). c, comparison of variant set meta-analysis *P* values from SMMAT and StocSum with the number of random vector replicates *B* being equal to 1,000. The x axis and the y axis represent −log_10_(*P*) from variant set meta-analysis using SMMAT and StocSum, respectively. The red line denotes the reference line of equality. Spearman’s rank correlation coefficients are shown at the bottom right. d, the Manhattan plot of variant set tests on LDL in the meta-analysis of ARIC AA and EA, and HCHS/SOL studies using StocSum. The x-axis represents the start physical chromosome and position of each variant set and the y-axis represents −log_10_(*P*) from the StocSum variant set test corresponding to SMMAT. The red line indicates the genome-wide significance level on the log scale, −log_10_ (3.7×10^−7^). All tests were performed on a high-performance computing server, with a single thread for each chromosome.

### LD score regression

StocSum can also be used to extend the LD Score Regression (LDSC) framework^14^ to underrepresented, admixed or isolated populations, without external reference panels. In this example, we compared LD scores and heritability estimates of four traits: LDL, high-density lipoprotein (HDL) cholesterol levels, systolic blood pressure (SBP), and diastolic blood pressure (DBP) from Hispanic/Latino adults in HCHS/SOL. LD scores were calculated using six different approaches: 1) StocSum (Sample): StocSum (*B*=1,000) on HCHS/SOL study samples; 2) LDSC (Sample): LDSC using HCHS/SOL study samples as internal reference panels; 3) LDSC (Eu): LDSC using European populations from the 1000 Genomes Project as external reference panels; 4) LDSC (EuAf): LDSC using European and African populations from the 1000 Genomes Project as external reference panels; 5) LDSC (EuAm): LDSC using European and American populations from the 1000 Genomes Project as external reference panels; and 6) LDSC (EuAfAm): LDSC using European, African and American populations from the 1000 Genomes Project as external reference panels. LD scores computed from StocSum (Sample) and LDSC using external reference panels were compared with those computed from LDSC (Sample).

LD scores from StocSum (Sample) were much closer to those from LDSC (Sample) (**Fig. 6e**), compared to LDSC results using external reference panels (**Fig. 6a-d**). Moreover, there seems to be an upward bias for many variants in LDSC (EuAf) and LDSC (EuAfAm) results, when African populations from the 1000 Genomes Project were included in the reference panel (**Fig. 6b,d**), highlighting the challenges in selecting appropriate external reference panels for LD score estimation in underrepresented, admixed or isolated populations. StocSum (Sample) required only 1.4% of CPU time used by LDSC (Sample) (**Fig. 6f**). It was also 6-fold to 42-fold faster than LDSC using different external reference panels.

**Figure 6:**
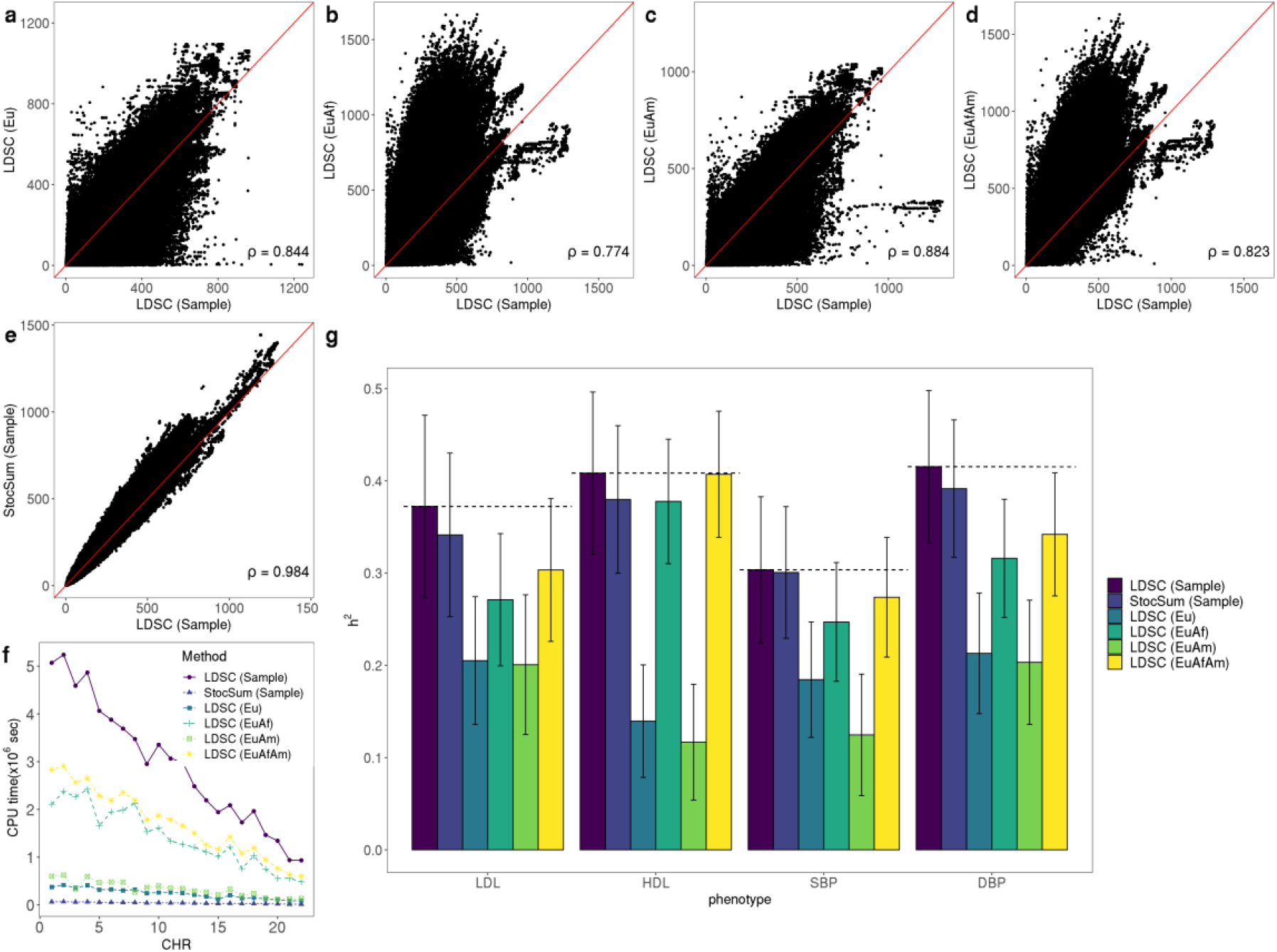
StocSum in LD score regression and heritability estimation. a-e, comparison of LD scores from LDSC (Sample) (x-axis) and different alternative methods (y-axis). a, LDSC (Eu). b, LDSC (EuAf). c, LDSC (EuAm). d, LDSC (EuAfAm). e, StocSum (Sample). Spearman’s rank correlation coefficients are shown at the bottom right. f, comparison of CPU time between StocSum and LDSC in LD score calculations. The x axis represents the chromosome numbers and the y axis represents the CPU time in 10^6^ seconds. g, heritability estimates using LD scores from LDSC and StocSum. The error bars show point estimates ± standard errors. LD scores were estimated from LDSC (Sample) and StocSum (Sample) using HCHS/SOL study samples, or LDSC on external reference panels using European, African and/or American populations from the 1000 Genomes Project: LDSC (Eu), LDSC (EuAf), LDSC (EuAm), and LDSC (EuAfAm).

Using LD scores from these six approaches, we compared heritability estimates of four traits LDL, HDL, SBP and DBP in HCHS/SOL (**Fig. 6g**). StocSum (Sample) results were consistently observed to be close to LDSC (Sample) heritability estimates, for all these traits. Heritability estimates from LDSC using external reference panels tended to be lower than LDSC (Sample), especially when African populations from the 1000 Genomes Project were excluded in the reference panel. For example, heritability estimates from LDSC (EuAm) were about 46.1%, 71.4%, 59.0%, and 51.1% lower compared to those from LDSC (Sample), for LDL, HDL, SBP, and DBP traits. Heritability estimates partitioned by different MAF bins also showed that StocSum (Sample) results were consistent with those from LDSC (Sample) (Fig. S14). Overall, StocSum is better suited for conducting LD score regression in Hispanic/Latino adults, while LDSC needs a reference panel that matches the LD structure in the study samples.

## Discussion

We have developed and implemented StocSum, a novel framework for generating, managing, and using stochastic summary statistics for WGS studies. We showed that in all the example applications that use between-variant correlation or LD matrices, either from the study samples or external reference panels, such as conditional association tests, variant set tests and LD score regression, we could use a much smaller stochastic summary statistic matrix ***U*** to replace the between-variant correlation or LD matrices, and flexibly extract the pairwise LD information between any variants on the same chromosome. This strategy was highly accurate and computationally efficient. The size of ***U*** scales linearly with the number of genetic variants *M*, compared to quadratically in the form of traditional pairwise LD matrices. The computing time for the stochastic summary statistic matrix ***U*** always scales linearly with both the sample size *N* and the number of genetic variants *M* (the same complexity with reading the data), regardless of any complex sample correlation structures. This matrix only needs to be computed once for each phenotype in both cross-sectional and longitudinal studies, and can be reused in single-variant tests, conditional association tests, and variant set tests with different variant set definitions.

StocSum leverages stochastic algorithms to reduce the computational burden in WGS studies. Similar algorithms have previously been applied to principal component analysis^77,78^, heritability^79^ and genetic correlation estimation^80^, and it is our hope that the StocSum framework can be extended to a wide range of other applications to genomic summary statistics that currently require external reference panels, thus facilitating use of genomic summary statistics from WGS studies. This is especially important for underrepresented, admixed, and/or isolated populations, for which appropriate reference panels are difficult to find. We have shown for variant set tests (Fig. S9–10) and LD score regression (Fig. 6) that external reference panels did not perform well even when all three ancestry populations for Hispanic/Latino adults were included, and the performance was even worse when a European-only reference panel was used. By using StocSum instead of external reference panels, more genetic research can be conducted in diverse populations that will equally benefit all humans.

StocSum will likely also facilitate international collaborations on genomic epidemiological research using WGS data, so that meta-analysis for rare genetic variants can be easily conducted without sharing individual-level WGS data across borders. Such collaborations have largely focused on common genetic variants in the past, by sharing genomic summary statistics. With the decreasing cost and increasing availability of WGS data, large-scale meta-analysis efforts on rare genetic variants are currently very difficult to coordinate, as variant sets determining how rare genetic variants should be grouped need to be pre-defined. In contrast, in the StocSum framework, researchers can combine the stochastic summary statistic matrices U from different studies first, and then decide how the variants should be grouped. When analysis plans change, there is no need to rerun any analyses using individual-level data, thus encouraging use of WGS data in international consortia.

WGS data are big in size and often difficult to share. Although large-scale studies such as the UK Biobank^11^, the TOPMed program^10^, and the CCDG initiative, have made plans to host their WGS data on cloud-computing platforms to facilitate access, it is still computationally expensive to directly analyzing individual-level data, making it financially difficult for small research groups to contribute to scientific discoveries using WGS data. The StocSum framework will democratize access to WGS resources, as we expect these high-level summary data will be generated by central analysis centers who are familiar with and have direct access to individual-level phenotype and WGS data, and broadly shared with the scientific community. All downstream analyses using StocSum are free of the sample size N and could be performed on a laptop. It is also an eco-friendly strategy by avoiding different research teams running individual-level WGS data analyses on the same phenotypes, which are at least *O*(*NM*) operations for each team, thus saving a lot of electricity in computation.

There are also several limitations. We have demonstrated concordance of StocSum results as compared to methods that directly use individual-level data, for both common and rare variants, but it does not imply these results are statistically valid in all scenarios. For example, asymptotic *P* values from GMMAT may not be well-calibrated for extremely unbalanced cases:control ratios from Biobank studies^81^. This issue likely also exists for StocSum tests, given the concordance of StocSum and GMMAT results, and would require further adjustments or approximations. Moreover, although LD scores and heritability estimates from StocSum matched well with those from LDSC using internal reference panels (Fig. 6), these heritability estimates are likely underestimates and may not compare with estimates from other studies, due to the relatively small sample size in HCHS/SOL. Also, the choice of the number of random vector replicates *B* depends on the scientific questions to be investigated in downstream analyses. It does not depend on the sample size *N*, although we note that for small studies with *N* < *B*, it might be more computationally expensive to use StocSum, compared to directly using individual-level data. In this study we have recommended using *B*=1,000 in all applications, and it worked well in variant set tests for both regions with the number of variants *q* ≤ *B* and *q* > *B* (Fig. S8). However, when it is of interest to test a very wide region with *q* being much greater than *B*, such as topologically associating domains ande chromosome-wide association by class of histone markers^82^, the performance of StocSum is not guaranteed. Nevertheless, we expect StocSum to be a computationally efficient and eco-friendly framework for WGS studies that will facilitate genetic research in diverse populations, international collaborations, and equal access to WGS resources for the scientific community.

## Methods

### Stochastic summary statistics

We first define the basic null model in the StocSum framework. Under the null hypothesis of no genetic fixed effects *H*_0_: ***β*** = 0, model (Eq.(1)) (see Results) reduces to

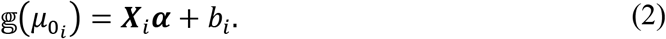

Here 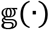 is a monotonic link function of *μ*_0_*i*__, and *μ*_0_*i*__ = *E*(*y_i_*|***X***_*i*_, *b_i_*) is the conditional mean of the phenotype *y_i_* under the null hypothesis *H*_0_: ***β*** = 0, given *p* covariates ***X***_*i*_ (including an intercept) and random effects *b_i_*, for individual *i* of *N* samples. Let 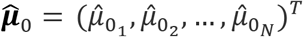 be a length *N* column vector for the estimated values of *μ*_0_*i*__, 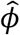 be an estimate of the dispersion parameter (or the residual variance for continuous traits in linear mixed models) *ϕ*, and 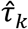 be the estimates for variance component parameters *τ_k_* corresponding to *N* × *N* relatedness matrices ***Φ***_*k*_, from the null model (Eq.(2)), we define 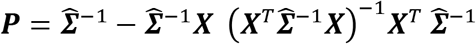 as the projection matrix, where 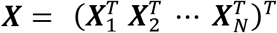 is a *N* × *p* covariate matrix, and 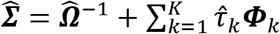 with 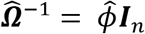 for continuous traits in linear mixed models, and 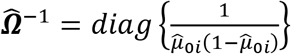 for binary traits in logistic mixed models^55^.

StocSum leverages a length *N* random vector ***R***_*b*_ from a multivariate normal distribution with mean **0** and covariance matrix ***P***, repeats this simulation process *B* times and combines ***R***_*b*_ (1 ≤ *b* ≤ *B*) into an *N* × *B* random matrix ***R*** = (***R***_1_ ***R***_2_ ⋯ ***R***_*B*_). In our implementation, we first decompose relatedness matrices ***Φ***_*k*_ = ***Z***_*k*_***Z***_*k*_^*T*^, where ***Z**_k_* is an *N* × *L_k_* matrix (*L_k_* ≤ *N*). For low-rank relatedness matrices (such as those indicating observations from the same sample in longitudinal studies), ***Z***_*k*_ is often known as the random effect design matrix, with *L_k_* being the rank of ***Φ***_*k*_. For sparse block-diagonal relatedness matrices (such as positive definite kinship matrices), ***Z***_*k*_ is the Cholesky decomposition of ***Φ***_*k*_, which is also sparse block-diagonal. We construct the *N* × *B* random matrix as 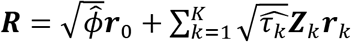, in which ***r***_0_ is an *N* × *B* random matrix and ***r***_*k*_ (1 ≤ *k* ≤ *K*) are *L_k_* × *B* random matrices, with all entries in ***r***_0_ and ***r***_*k*_ simulated from a standard normal distribution.

For an *N* × *M* genotype matrix ***G*** for *M* variants on the whole genome (or on one chromosome), the *M* × *B* stochastic summary statistic matrix ***U*** can be calculated as ***U*** = ***G***^*T*^***R***. In the next sections, we describe how the stochastic summary statistics can be used in various downstream genetic analysis applications.

### Single-variant tests

We are interested in conducting single-variant tests for the null hypothesis *H*_0_: *β* = 0, using the score test. The GMMAT single-variant score is 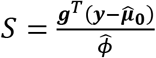, where ***g*** = (*g*_1_ *g*_2_ ⋯ *g_N_*)^*T*^ is a length *N* column genotype vector for the variant of interest, ***y*** = *y*_1_ *y*_2_ ⋯ *y_N_*)^*T*^ is a length *N* column vector for the phenotype (Chen et al., 2016). The variance of the score is *Var*(*S*|*H*_0_) = ***g***^*T*^***Pg***.

Denote the *j*th row of the stochastic summary statistic matrix ***U*** (for variant *j*, 1 ≤ *j* ≤ *M*) by a length *B* row vector ***U***_*j*_, we can show that the variance *Var*(*S*|*H*_0_) of single-variant score *S* for variant *j* can be estimated as 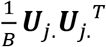, without using any individual-level data. The asymptotic *P* value is then computed using the single-variant score *S*^55^ and its variance estimated from the stochastic summary statistic matrix ***U***, for each variant of interest.

### Conditional association tests

Assume *Ġ* is an *N* × *c* genotype matrix for *c* ≥ 1 association genetic variants to be conditioned on and ***g*** is a length *N* column genotype vector for the variant of interest in the conditional association test. The single-variant score conditional on the variant set ***Ġ*** is

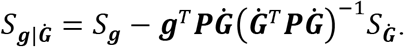

The variance of the conditional score is *Var*(*S*_***g|Ġ***_) = ***g***^*T*^***Pg*** – ***g***^*T*^***PĠ***(***Ġ***^*T*^***PĠ***)^−1^***Ġ***^*T*^***Pg***^17^.

In the StocSum framework, *S_**g**_* and ***U_g_*** are the single-variant score and stochastic summary statistics corresponding to the variant of interest in the conditional association test and ***S_Ġ_*** (a length *c* vector) and ***U_Ġ_*** (a *c* × *B* matrix) are the single-variant score and stochastic summary statistics corresponding to the association variants to be conditioned on. The conditional score can be computed as

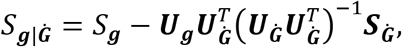

and the conditional stochastic summary statistics can be computed as

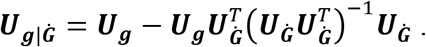

The variance *Var*(*S*_***g|Ġ***_) of the conditional score *S*_***g|Ġ***_ can be estimated as 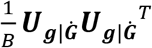. The asymptotic *P* value is computed using the conditional score *S*_***g|Ġ***_ and its variance estimated from the stochastic summary statistics ***U*_*g|Ġ*_**, for each variant of interest in the conditional association test.

### Gene-environment interaction tests

We introduce a general model for testing *m* gene-environment interaction (GEI) terms in the GLMM framework. The full model including the genetic main effect and GEI effects is

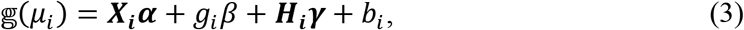

where *g_i_* is the genotype for the variant of interest for individual *i*, *β* is a scalar of the genetic main effect, ***H***_*i*_ is a length *m* row vector for the GEI terms, which include the products of *g_i_* and *m* environmental factors (a subset from *p* covariates in ***X***_*i*_), and ***γ*** is a length *m* column vector for GEI effects. We note that under the constraint ***γ*** = 0, *β* also represents the marginal genetic effect. Other notations follow the null model (Eq.(2)).

The single-variant score for the marginal genetic effect is 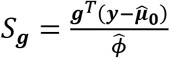 and its variance is *Var*(*S_**g**_*) = ***g***^*T*^***Pg***. The single-variant score for the GEI effects is 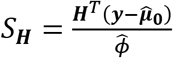 and its *m* × *m* covariance matrix is *Var*(*S_**H**_*) = ***H***^*T*^***PH***, where 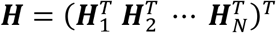 is a *N* × *m* matrix for the GEI terms. The score for GEI effects adjusting for the marginal genetic effect can be approximated by *S_***H|g***_* = *S_**H**_* – ***H***^*T*^***Pg***(***g***^*T*^***Pg***)^−1^*S_**g**_*^61^, with a covariance matrix *Var*(*S_***H|g***_*) = ***H***^*T*^***PH*** – ***H***^*T*^***Pg***(***g***^*T*^***Pg***)^−1^***g***^*T*^***PH***. The marginal genetic effect can be tested using the quadratic form *S****_g_***^*T*^*Var*(*S_**g**_*)^−1^*S_**g**_*, which follows a chi-square distribution with 1 degree of freedom under the null hypothesis of no marginal genetic effects. The GEI effects can be tested using *S*_***H|g***_^*T*^*Var*(*S_***H|g***_*)^−1^*S_***H|g***_*, which follows a chi-square distribution with *m* degrees of freedom under the null hypothesis of no gene-environment interactions. The joint test, which evaluates both marginal genetic effects and GEI effects, can be constructed by the sum of these two chi-square statistics, since *S_**H**_* and *S_**H|g**_* are asymptotically independent. The joint test statistic follows a chi-square distribution with 1 + *m* degrees of freedom under the null hypothesis of no marginal genetic effects or gene-environment interactions.

In the StocSum framework, we first compute stochastic summary statistics for the marginal genetic effect ***U_g_*** = ***g***^*T*^***R*** and GEI effects ***U_H_*** = ***H***^*T*^***R*** using individual-level data. We can use 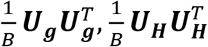, and 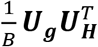 to estimate the variance of the marginal genetic effect score *Var*(*S_**g**_*), the covariance matrix of the GEI effect score *Var*(*S_**H**_*), and the covariance of *S_**g**_* and *S_**H**_*, respectively. The adjusted scores can be constructed as 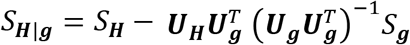, and its variance *Var*(*S_***H|g***_*) can be approximated as 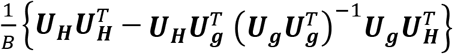.

### Variant set tests

We include four variant set tests: the burden test^34–37^, SKAT^38^, SKAT-O^83^, and the efficient hybrid test of burden and SKAT^21,39^, in the StocSum framework. Here we consider a variant set including *q* genetic variants (*q* > 1) and denote 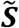 as a length *q* single-variant score vector, and 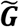 as an *N* × *q* genotype matrix (a subset of the *N* × *M* genotype matrix ***G*** on the whole genome, or on one chromosome). We note that our examples are not a complete list of all variant set tests that are commonly used, but any other variant set tests that would require *q* × *q* covariance matrices could also be implemented using stochastic summary statistics.

The burden test statistic can be constructed as

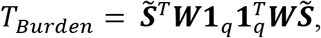

where ***W*** = *diag*{*w_j_*} is a pre-specified *q* × *q* diagonal weight matrix, and **1**_*q*_ is a length *q* vector of 1’s. The weights can be a function of the MAF^36,38^, or functional annotation scores such as CADD^84,85^, FATHMM-XF^86^, and annotation principal components from STAAR^87^. Under the null hypothesis, the statistic *T_Burden_* asymptotically follows 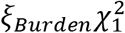, where the scaling factor 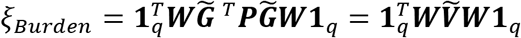 (where 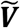 is a *q* × *q* covariance matrix for the single-variant score vector 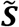), and 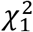 is a chi-square distribution with 1 df. In the StocSum framework, *ξ_Burden_* can be estimated as 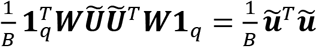, where 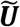 is a *q* × *B* matrix (a subset of the *M* × *B* stochastic summary statistic matrix ***U***), and 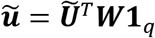 is a length *B* vector (i.e., column sum of 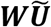).

The SKAT statistic can be constructed as

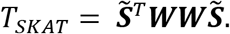

Under the null hypothesis, *T_SKAT_* asymptotically follows 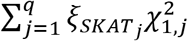, where 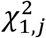 are independent chi-square distributions with 1 df, and *ξ_SKAT_j__* are the eigenvalues of 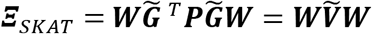. In the StocSum framework, *ξ_SKAT_j__* can be estimated as the square of the singular values of 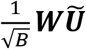 (**Supplementary Note 1**).

In SKAT-O, the variance component statistic *T_ρ_* given a weight parameter *ρ* (0 ≤ *ρ* ≤ 1) is

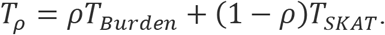

If *ρ* = 1, *T_ρ_* becomes the burden test statistic *T_Burden_*; if *ρ* = 0, *T_ρ_* becomes the SKAT statistic *T_SKAT_*. SKAT-O searches for an optimal *ρ* by minimizing the *P* value of *T_ρ_*. Specifically, the *q* × *q* weighted covariance matrix 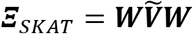 is decomposed into two parts 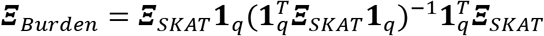 and ***Ξ**_SKAT|Burden_* = ***Ξ**_SKAT_* – ***Ξ**_Burden_*, used in subsequent one-dimensional numerical integration to compute the SKAT-O *P* value. In the StocSum framework, ***Ξ**_Burden_* can be estimated as 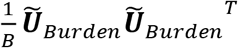, where 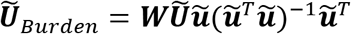, and ***Ξ**_SKAT|Burden_* can be estimated as 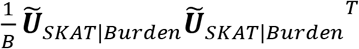, where 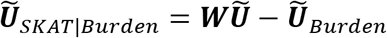.

In the efficient hybrid test to combine the burden test and SKAT, the adjusted SKAT statistic *T_SKAT|Burden_* can be approximated by

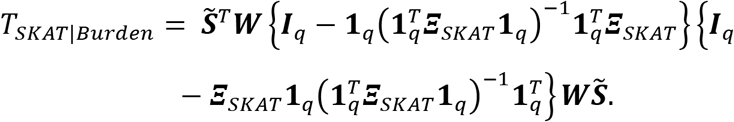

Under the null hypothesis, *T_SKAT|Burden_* asymptotically follows 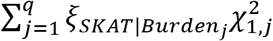, where 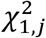 are independent chi-square distributions with 1 df and *ξ_SKAT|Burden_j__* are the eigenvalues of ***Ξ**_SKAT|Burden_*. In the StocSum framework, these eigenvalues can be estimated as the square of the singular values of 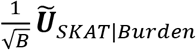 (**Supplementary Note 2**).

### Meta-analysis

In a traditional meta-analysis on a region with *q* genetic variants from *L* studies, we use the single-variant scores 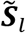 and the covariance matrix 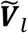 from each study *l* (1 ≤ *l* ≤ *L*). The variant set meta-analysis can be performed using the summary scores 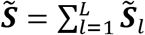 and the summary covariance matrix 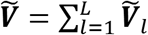^18,19,21,31,33^. The single-variant meta-analysis only requires 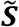 and the diagonal elements of 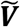. In the StocSum framework, we compute 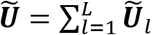 instead of 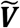. Assuming *q* < *B*, each column of 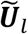 follows a multivariate normal distribution with mean **0** and covariance matrix 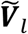, and 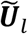 are independent across *L* studies assuming no sample overlaps or between-study relatedness. Therefore, each column of 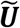 follows a multivariate normal distribution with mean **0** and covariance matrix 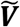. In our implementation, we first compute the stochastic summary statistic matrix 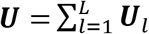 for all *M* genetic variants on the whole genome (or one chromosome), regardless of how variants should be grouped, and then extract *q* genetic variants by taking a subset of ***U*** only when computing *P* values, for both single-variant meta-analysis and variant set meta-analysis.

### LD score regression

LD Score Regression (LDSC) has been widely applied to GWAS summary statistics to estimate confounding bias, heritability explained by genotyped variants, heritability enrichments of functional categories, and genetic correlations^14,15,88^. The classical LDSC model can be written as

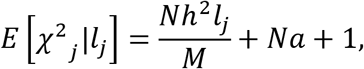

where *χ*^2^*j* denotes the *χ*^2^ statistic of variant *j* from GWAS summary statistics; 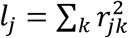 is the LD score of variant *j* with 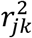 being the squared Pearson correlation coefficient of genotypes between variants *j* and *k*, *N* is the sample size, *M* is the total number of variants, *a* is a measure of confounding bias, and *h*^2^ is the heritability of the phenotype. In practice, LDSC calculates *l_j_* by summing up 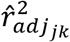 for all variants *k* in specific window around the index variant *j*. The adjusted correlation estimate 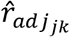 can be computed from the sample correlation estimate 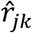 using

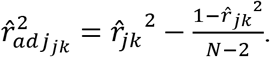

Sample correlation coefficients 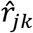 can be estimated as 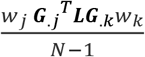, where *G_.j_* is the *j*th column of the genotype matrix ***G***, representing variant *j*, ***L*** = (***I***_*N*_ – **1**_*N*_(**1**_*N*_^*T*^**1**_*N*_)^−1^**1**_*N*_^*T*^) is an *N* × *N* idempotent projection matrix, and 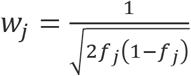 (*f_j_* is the MAF of variant *j*) is a weight that standardizes *G_.j_* to a unit variance.

In the StocSum framework, we construct the *N* × *B* random matrix as ***R = Lr***_0_, where ***r***_0_ is an *N* × *B* random matrix with all entries simulated from a standard normal distribution. For an *N* × *M* genotype matrix ***G*** for all *M* genetic variants on the whole genome (or one chromosome), we compute the *M* × *B* stochastic summary statistic matrix ***U*** = ***WG***^*T*^***R***, where ***W*** = *diag*{*w_j_*} is an *M* × *M* diagonal weight matrix. For variant *j*, we subset *M_j_* variants within the flanking region (with a default window width of 1000 Kb) to get the corresponding *M_j_* × *B* subset 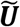. The adjusted correlation coefficient 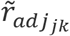 for 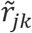 from StocSum is computed as (**Supplementary Note 3**)

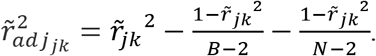

The LD score *l_j_* of variant *j* could be estimated by summarizing stochastic summary statistics of *M_j_* variants in flanking region,

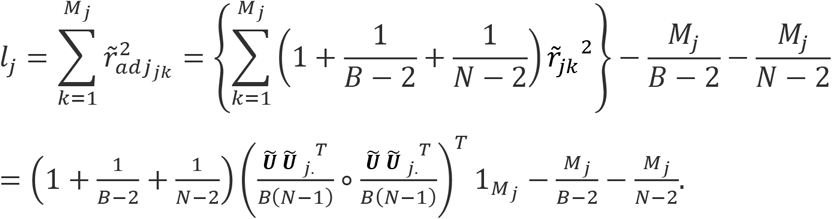

in which ∘ denotes the Hadamard product, and 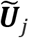. is the *j*th row of 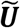.

### Whole genome sequence and phenotype data

The Trans-Omics for Precision Medicine (TOPMed), sponsored by the National Heart, Lung and Blood Institute (NHLBI), generates scientific resources to enhance our understanding of fundamental biological processes that underlie heart, lung, blood and sleep disorders (HLBS)^10^. WGS of the TOPMed samples was performed over multiple studies, years and sequencing centers. The TOPMed freeze 8 WGS data include 138K samples from 72 studies. The sequence reads were aligned to the human genome build GRCh38 using BWA-MEM following the protocol published previously^89^. To perform variant quality control, a support vector machine classifier was trained on known variant sites (positive labels) and Mendelian inconsistent variants (negative labels). Further variant filtering was done for variants with excess heterozygosity and Mendelian discordance. Sample quality control measures included: concordance between annotated and inferred genetic sex, concordance between prior array genotype data and TOPMed WGS data, and pedigree checks^10^.

In this paper, our analysis includes genotypes and phenotypes from two TOPMed studies, Hispanic Community Health Study/Study of Latins (HCHS/SOL) and the Atherosclerosis Risk in Communities (ARIC) study.

#### HCHS/SOL data

The HCHS/SOL is a multi-center study of Hispanic/Latino populations with the goal of determining the role of acculturation in the prevalence and development of diseases, and to identify other traits that impact Hispanic/Latino health^90^. Participants were recruited using a multi-stage probability sample design, as described previously^90,91^. The HCHS/SOL is composed of six different background groups including Central Americans, Cubans, Dominicans, Mexicans, Puerto Ricans, and South Americans^7^. A total of 123,004,674 variants from 7,684 HCHS/SOL participants in TOPMed were available for genetic association analyses.

Low-density lipoprotein (LDL) cholesterol levels were used as an illustrating example in single-variant tests, conditional association tests, variant set tests, meta-analysis, and LD score regression. Additional phenotypes including high-density lipoprotein (HDL) cholesterol levels, systolic blood pressure (SBP), and diastolic blood pressure (DBP) were also used as examples in LD score regression. To account for the effect of lipid-lowering medication, LDL cholesterol levels for study participants who took statins were adjusted by dividing raw values by 0.7, following previous studies^57,92,93^. Both LDL and HDL cholesterol levels were set to missing for study participants with unknown statins use, unknown fibric/nicotinic acids use, or those who took only fibric/nicotinic acids but no statins. SBP and DBP were adjusted by adding 15 mmHg and 10 mmHg for study participants self-reporting use of any antihypertensive medication, respectively^76^. The waist-hip ratio (WHR) was used as an illustrating example in gene-environment interaction tests.

#### ARIC data

The cohort component of the ARIC study began in 1987, and each of the four ARIC field centers (Washington County, MD; Forsyth County, NC; Jackson, MS; and Minneapolis, MN) randomly selected and recruited a cohort sample of approximately 4,000 individuals aged 45-64 from a defined population in their community. A total of 15,792 participants received an extensive examination, including medical, social, and demographic data^94^. These participants were examined with the first (baseline) exam occurring in 1987-89, the second in 1990-92, the third in 1993-95, the fourth in 1996-98, the fifth in 2011-13, and the sixth in 2016-17. The TOPMed WGS study over-sampled ARIC participants with incident venous thromboembolism (VTE). We removed samples/visits with missing phenotype (LDL) or covariates (age, sex, BMI, field center, and top five ancestry principal components), resulting in 26,668 observations from 6,327 ARIC EA samples and 7,514 observations from 2,045 ARIC AA samples. After removing low-quality variants with a genotype call rate less than 90% and monomorphic markers, there were 91,715,717 and 69,958,574 variants in ARIC EA and AA samples, respectively.

Longitudinal LDL cholesterol levels from the baseline exam until up to the 6th exam were used as an illustrating example in single-variant and variant set meta-analyses. To account for the effect of lipid-lowering medication, LDL cholesterol levels for study participants who took statins were adjusted by dividing raw values by 0.7^57,92,93^. LDL cholesterol levels were set to missing for study participants with unknown statins use, unknown cholesterol medication use, or inconsistent information from statins use and cholesterol medication use.

#### Reference data from 1000 Genomes

Individual-level WGS data from the 1000 Genomes Project^95^ were used as reference panels in fastBAT variant set tests and LD score regression. Only high-quality variants with a genotype call rate ≥ 95% and passed the quality control filters were included. Four reference panels were constructed with different combinations of super-populations: European (Eu), European and African (EuAf), European and American (EuAm), and European, African and American (EuAfAm), with 23,654,568, 45,780,202, 31,334,904, and 49,350,7868 variants from 503, 894, 682, and 1,073 samples, respectively.

### Statistical Analyses

#### Single-variant tests

We removed samples with missing values in the phenotype LDL cholesterol levels or covariates (age, sex, body mass index [BMI], field center, sampling weight, Hispanic/Latino background groups, and top five ancestry principal components) and excluded variants with a genotype call rate less than 90% and monomorphic markers in single-variant test comparisons. After quality control, a total of 120,066,450 variants from 7,297 HCHS/SOL samples were available for analysis. We included age, age^2^, sex, age × sex, age^2^ × sex, BMI, field center, sampling weight, Hispanic/Latino background groups and top five ancestry principal components as fixed-effects covariates. We rank-normalized residuals after regressing the phenotype LDL cholesterol levels on fixed-effects covariates, and then used them as the phenotype in downstream null model fitting and association tests^96^. Three random effects representing household, census block, and kinship effects were included to account for sample relatedness. We also allowed the residual variance to be different across 6 Hispanic/Latino background groups (i.e., Central American, Cuban, Dominican, Mexican, Puerto Rican, and South American), in a heteroscedastic linear mixed model^7^ for both GMMAT and StocSum. The *P* values from StocSum were compared to those from GMMAT using individual-level data. The default value of the number of random vectors *B* in StocSum was set to 1,000. To benchmark the numerical accuracy and required computational resources, the number of random vectors *B* changed from 10 (StocSum (B=10)), 100 (StocSum (B=100)), 1,000 (StocSum (B=1,000)), to 10,000 (StocSum (B=10,000)).

To compare with fastGWA^60^ in single-variant analysis, we dropped household and census block random effects, and only included a kinship random effects to account for sample relatedness. We also assumed an equal residual variance across 6 Hispanic/Latino background groups in the linear mixed model for GMMAT and StocSum, to make a fair comparison with fastGWA.

#### Conditional association tests

We performed conditional analyses for the seven regions associated with LDL at the genome-wide significance level of 5×10^−8^ from the single-variant analysis in HCHS/SOL (**Table S1**). For each region, we started with a sentinel variant with the smallest *P* value, and computed conditional association test *P* values for all variants in the flanking region (1 Mb) after adjusting for the sentinel variant. If there were any variants in a region with a conditional *P* < 5×10^−8^, we then selected the variant with the smallest conditional *P* value as the secondary association variant, and performed conditional analyses after adjusting for both association variants.

#### Gene-environment interaction tests

We compared gene-environment interaction tests in StocSum with MAGEE single-variant interaction tests using individual-level data. We focused on gene-sex interaction effects on an anthropometric phenotype waist-hip ratio (WHR) which shows strong evidence of sex dimorphism, using WGS data from HCHS/SOL. We included age, age^2^, sex, age × sex, age^2^ × sex, BMI, field center, sampling weight, Hispanic/Latino background groups, top five ancestry principal components (PCs), and sex by top five ancestry PC interactions as fixed-effects covariates. After removing samples with missing values in the phenotype WHR or covariates, and variants with a genotype call rate less than 90% and monomorphic markers, a total of 122,076,760 variants from 7,636 HCHS/SOL samples were available for analysis. Similar to the single-variant analysis, we followed a two-step approach^96^ and used rank-normalized WHR residuals as the phenotype in null model fitting and gene-sex interaction tests. We included three random effects representing household, census block and kinship effects to account for sample relatedness, and used a heteroscedastic linear mixed model by allowing the residual variance to be different across the 12 sex by Hispanic/Latino background groups. The marginal genetic effect, gene-sex interaction, and joint test *P* values from StocSum were compared to corresponding test results from MAGEE single-variant interaction tests using individual-level data.

#### Variant set tests

We compared variant set tests using StocSum versus SMMAT using individual-level data. After removing samples with missing values in the phenotype LDL cholesterol levels or covariates, and variants with a genotype call rate less than 90% and monomorphic markers, a total of 120,066,450 variants from 7,297 HCHS/SOL samples were available for analysis. We used the same null model as previously described in the single-variant tests for GMMAT and StocSum, and conducted a sliding window analysis^43^ with 20kb non-overlapping windows. We applied a beta density function with parameters 1 and 25 on the MAF as variant weights^38^ in both SMMAT and StocSum. SMMAT requires individual-level data to conduct variant set tests. In contrast, StocSum directly uses the single-variant summary statistics and stochastic summary statistics previously computed for single-variant tests.

To compare with fastBAT^97^ in variant set tests, we used the same kinship-only null model with equal residual variance as previously described in the single-variant test comparison for fastGWA, GMMAT, and StocSum. We also changed variant weights using a beta density function with parameters 0.5 and 0.5 on the MAF (also known as the Madsen-Browning weights)^98^, equivalent to rescaling the genotypes to a unit variance in fastBAT. Four external reference panels from 1000 Genomes (Eu, EuAf, EuAm, EuAfAm), as well as an internal reference panel using the HCHS/SOL study samples, were used to estimate LD between variants in each set in fastBAT.

In a second example, we also applied StocSum to variant set tests using windows defined by functional genomic units. We collected Hi-C data generated from an *in situ* Hi-C protocol on human GM12878 B-lymphoblastoid cells^49^, in which the crosslinked DNA was pulled down followed by Illumina sequencing. The whole genome was split into non-overlapping segments with a bin size of 10kb (i.e., contact matrices were generated at base pair delimited resolutions of 10kb), and a total of 17,224 pairs of contacts were defined. Each segment pair can be considered as a long-distance DNA crosslink. We grouped variants from each contact pair as a variant set, including two 10kb windows which may not be located in close proximity on the primary structure of DNA (the linear sequence), to evaluate the performance of StocSum on variant sets that are physically farther away and not typically covered using fixed-size sliding windows.

#### Meta-analysis

We combined StocSum on LDL cholesterol levels from ARIC and HCHS/SOL in single-variant and variant set meta-analysis. For HCHS/SOL, we used single-variant summary statistics and stochastic summary statistics previously computed for single-variant tests on LDL cholesterol levels. For ARIC, we first fit two linear mixed models separately for EA and AA, treating LDL cholesterol levels from up to 6 visits as repeated measures for each participant, and then computed single-variant summary statistics and stochastic summary statistics. We included age, age^2^, sex, age × sex, age^2^ × sex, BMI, field center, and top five ancestry principal components as fixed-effects covariates. We rank-normalized residuals after regressing the phenotype LDL cholesterol levels on fixed-effects covariates, and then used them as the phenotype^96^. In each ARIC dataset (EA and AA), variants with a genotype call rate less than 90% and monomorphic markers were excluded. After quality control, there were a total of 91,715,717 variants from 6,327 ARIC EA samples, and 69,958,574 variants from 2,045 ARIC AA samples.

We took the union of all variants and combined ARIC EA, ARIC AA, and HCHS/SOL summary statistics in a traditional single-variant meta-analysis using GMMAT, and a traditional variant set meta-analysis using SMMAT. In StocSum meta-analysis, we combined stochastic summary statistics from ARIC EA, ARIC AA and HCHS/SOL into a single file by adding together stochastic summary statistics for the same variant across three studies. We assigned 0 to both the single-variant summary statistic and stochastic summary statistic for a variant that was not observed in a study, since it did not contribute to the test statistic. In variant set meta-analysis, we applied a beta density function with parameters 1 and 25 on the MAF as variant weights, and conducted a 20kb sliding window analysis, for both SMMAT and StocSum.

#### LD score regression

In LD score regression, we only included common genetic variants with MAF ≥ 1% in HCHS/SOL. Following previous guidelines^14,16,99,100^, we excluded variants within the major histocompatibility complex (MHC; chromosome 6: 25-34Mb) and variants in regions with exceptionally long-range LD (**Table S6**). After quality control, 11,190,311 common variants with MAF > 1% from 7,289 HCHS/SOL study samples were used in StocSum to calculate LD score. We used single-variant summary statistics from GWAS of LDL, HDL, SBP and DBP in HCHS/SOL using GMMAT. Covariates included age, age^2^, sex, age × sex, age^2^ × sex, BMI, field center, sampling weight, Hispanic/Latino background groups, and top five ancestry principal components. The same HCHS/SOL study samples were used as an internal reference panel in the LDSC program and StocSum to calculate LD scores, i.e., LDSC (Sample) and StocSum (Sample). With the same filters, four external reference panels from the 1000 Genomes Project were used in the LDSC program to calculate LD scores, i.e., LDSC (Eu), LDSC (EuAf), LDSC (EuAm), LDSC (EuAfAm), including 9,092,238, 14,296,986, 9,410,628, 13,819,023 common variants with MAF > 1% (1000 Genomes Project Consortium), from 503 Eu, 894 EuAf, 682 EuAm, and 1,073 EuAfAm samples, respectively. With the LD scores from these internal and external references, the LDSC program was used to estimate heritability. For both LDSC and StocSum, we used a 1 Mb window around each index variant to calculate its LD score.

To evaluate the performance of StocSum, we also compared heritability estimates from LDSC (Sample) and StocSum (Sample) partitioned by different MAF bins. Common variants from HCHS/SOL and external reference panels were divided into 6 MAF bins, i.e., 1% < MAF ≤ 5%, 5% < MAF ≤ 10%, 10% < MAF ≤ 20%, 20% < MAF ≤ 30%, 30% < MAF ≤ 40%, and 40% < MAF ≤ 50%. Partitioned LD scores for different MAF bins were calculated by LDSC and StocSum, i.e., LDSC (Sample), LDSC (Eu), LDSC (EuAf), LDSC(EuAm), LDSC (EuAfAm), and StocSum (Sample). Partitioned heritability was estimated by the LDSC program with summary statistics for the phenotype LDL and partitioned LD scores.

## Acknowledgements

Molecular data for the Trans-Omics in Precision Medicine (TOPMed) program was supported by the National Heart, Lung and Blood Institute (NHLBI). Whole genome sequencing for “NHLBI TOPMed - NHGRI CCDG: Hispanic Community Health Study/Study of Latinos (HCHS/SOL) (phs001395.v1.p1)” was performed at Baylor College of Medicine Human Genome Sequencing Center (HHSN268201600033I). Whole genome sequencing for “NHLBI TOPMed - NHGRI CCDG: Atherosclerosis Risk in Communities (ARIC) (phs001211.v1.p1)” was performed at Baylor College of Medicine Human Genome Sequencing Center (3U54HG003273-12S2; HHSN268201500015C) and the Broad Institute Genomics Platform (3R01HL092577-06S1). Core support including centralized genomic read mapping and genotype calling, along with variant quality metrics and filtering were provided by the TOPMed Informatics Research Center (3R01HL-117626-02S1; contract HHSN268201800002I). Core support including phenotype harmonization, data management, sample-identity QC, and general program coordination were provided by the TOPMed Data Coordinating Center (R01HL-120393; U01HL-120393; contract HHSN268201800001I). We gratefully acknowledge the studies and participants who provided biological samples and data for TOPMed.

The Genome Sequencing Program (GSP) was funded by the National Human Genome Research Institute (NHGRI), the National Heart, Lung, and Blood Institute (NHLBI), and the National Eye Institute (NEI). The GSP Coordinating Center (U24 HG008956) contributed to cross program scientific initiatives and provided logistical and general study coordination. The Centers for Common Disease Genomics (CCDG) program was supported by NHGRI and NHLBI, and whole genome sequencing was performed at the Baylor College of Medicine Human Genome Sequencing Center (UM1 HG008898).

The Hispanic Community Health Study/Study of Latinos is a collaborative study supported by contracts from the National Heart, Lung, and Blood Institute (NHLBI) to the University of North Carolina (HHSN268201300001I / N01-HC-65233), University of Miami (HHSN268201300004I / N01-HC-65234), Albert Einstein College of Medicine (HHSN268201300002I / N01-HC-65235), University of Illinois at Chicago – HHSN268201300003I / N01-HC-65236 Northwestern Univ), and San Diego State University (HHSN268201300005I / N01-HC-65237). The following Institutes/Centers/Offices have contributed to the HCHS/SOL through a transfer of funds to the NHLBI: National Institute on Minority Health and Health Disparities, National Institute on Deafness and Other Communication Disorders, National Institute of Dental and Craniofacial Research, National Institute of Diabetes and Digestive and Kidney Diseases, National Institute of Neurological Disorders and Stroke, NIH Institution-Office of Dietary Supplements.

The Atherosclerosis Risk in Communities study has been funded in whole or in part with Federal funds from the National Heart, Lung, and Blood Institute, National Institutes of Health, Department of Health and Human Services, under Contract nos. (75N92022D00001, 75N92022D00002, 75N92022D00003, 75N92022D00004, 75N92022D00005). The authors thank the staff and participants of the ARIC study for their important contributions.

This work was supported by NHLBI grant R01 HL145025.

## Competing interests

The authors declare no competing interests.

## Supplementary Note

### 1. Approximating eigenvalues in variant set tests using singular values from StocSum

For *q* variants (*q* < *B*), the *q* × *q* covariance matrix used in variant set tests is 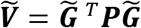. In the StocSum framework, we compute a *q* × *B* matrix 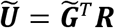, where each column ***R***_*b*_ (1 ≤ *b* ≤ *B*) of an *N* × *B* random matrix ***R*** = (***R***_1_ ***R***_2_ ⋯ ***R***_*B*_) is a length *N* random vector generated from a multivariate normal distribution with mean **0** and covariance matrix ***P***. Each column 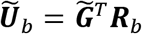 of 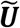 then follows a multivariate normal distribution with mean **0** and covariance matrix 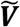, and the *B* columns of 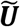 are independent and identically distributed. Therefore, when *B* is large, 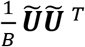 converges to the covariance matrix 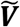. For 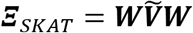 in SKAT, we can use 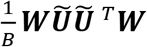 to estimate ***Ξ***_*SKAT*_.

We compute the singular value decomposition 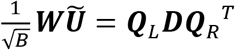, where *r* ≤ min(*q*, *B*) is the rank of 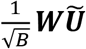, ***Q***_*L*_ and ***Q***_*R*_ are *q* × *r* and *B* × *r* semi-unitary matrices, respectively (***Q***_*L*_^*T*^***Q***_*L*_ = ***Q***_*R*_^*T*^***Q***_*R*_ = ***I***_*r*_), and ***D*** is an *r* × *r* diagonal matrix with elements being the singular values of 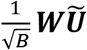. As we use 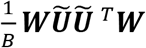 to estimate ***Ξ***_*SKAT*_, where 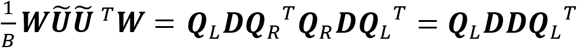, elements in the *r* × *r* diagonal matrix ***DD*** (the square of the singular values of 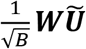) can be used to estimate the eigenvalues of ***Ξ***_*SKAT*_ when *r* = *q*. If *r* < *q* (for example, when testing a large genomic region with *q* > *B*), we could only estimate the top *r* (which is usually equal to *B* when *q* > *B*) eigenvalues of ***Ξ***_*SKAT*_ using the singular values of 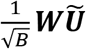.

### 2. Approximating eigenvalues in the efficient hybrid variant set test using singular values from StocSum

In the efficient hybrid variant set test to combine the burden test and SKAT, the adjusted SKAT statistic asymptotically follows a weighted sum of independent chi-square distributions with 1 df, where the weights are the eigenvalues of

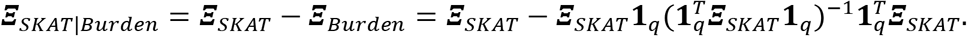

As we use 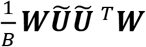 to estimate ***Ξ***_*SKAT*_ (**Supplementary Note 1**), let 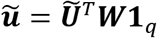 be a length *B* vector denoting the column sum of 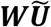, and define 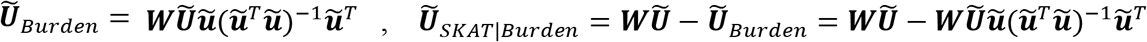 (see **Methods**), it follows that

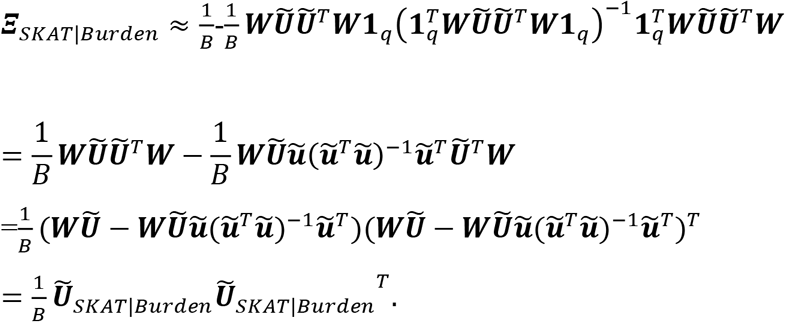

Therefore, similar to **Supplementary Note 1**, the eigenvalues of the *q* × *q* matrix ***Ξ***_*SKAT|Burden*_ can be estimated using the square of the single values of the *q* × *B* matrix 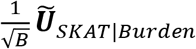.

### 3. Derivation of the adjusted correlation coefficient in the StocSum framework

Let *r_jk_* be the Pearson correlation coefficient between variants *j* and *k*, the sample correlation coefficient 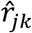 can be estimated using individual-level centered and rescaled genotypes (with mean 0 and variance 1), namely, 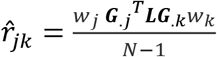, where ***G***_*.j*_ and ***G***_*.k*_ are the *j*th and *k*th columns of the full genotype matrix ***G***, representing variants *j* and *k*, *w_j_* and *w_k_* are rescaling weights that standardize genotypes to a unit variance, and ***L*** = (***I***_*N*_ – **1**_*N*_(**1**_*N*_^*T*^**1**_*N*_)^−1^**1**_*N*_^*T*^) is an *N* × *N* idempotent projection matrix that centers the genotypes (see **Methods**). The asymptotic distribution of 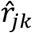 is given by

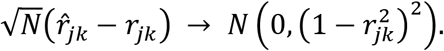

Therefore,

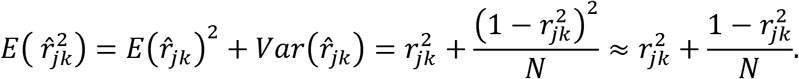

In LD score regression, the higher order term is ignored and the adjusted squared correlation coefficient is computed as 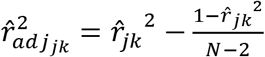 to reduce the bias (Bulik-Sullivan et al., 2015).

In the StocSum framework, we compute the *M* × *B* stochastic summary statistic matrix ***U*** = ***WG***^*T*^***R***, where ***W*** = *diag*{*w_j_*} is an *M* × *M* diagonal weight matrix, and ***G*** is an *N* × *M* genotype matrix for all *M* genetic variants on the whole genome (or one chromosome). We use ***U***_*j*_. and ***U***_*k*_. to denote length *B* row vectors from ***U*** for variants *j* and *k*, respectively. Then we can use 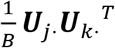 to estimate *w_j_* ***G***_*.j*_^*T*^***LG***_*.k*_*w_k_*, and therefore 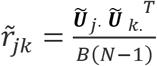 converges to 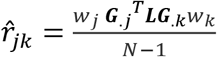 when *B* is large. Given 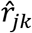, the asymptotic distribution of 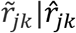 follows

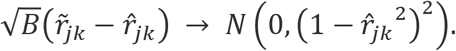

Therefore,

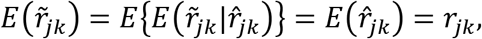

and ignoring the higher order terms in the variance, we have

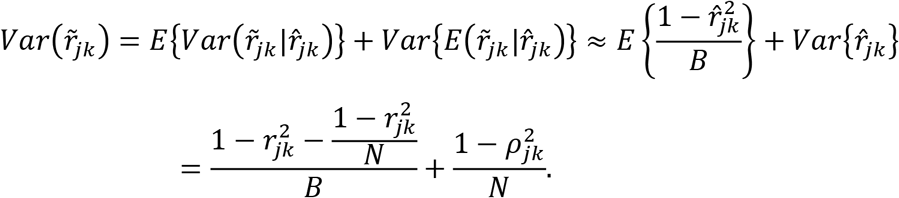

Hence,

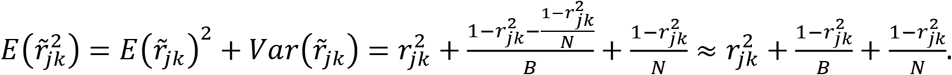

The term 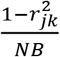 is ignored as both *N* and *B* are large. Following the same adjustment in LDSC (Bulik-Sullivan et al., 2015), we calculate adjusted correlation coefficient 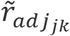 for 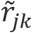 from StocSum using

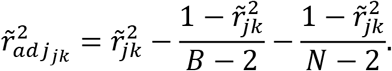

## Supplementary Tables

**Table S1.** Significant association regions with LDL cholesterol levels from single-variant tests in HCHS/SOL. Only variants with MAF > 0.5% were included. Genome coordinates presented were based on GRCh38.

**Table S2.** Regions showing suggestive evidence of gene-sex interactions or genetic associations accounting for gene-sex interactions on WHR in HCHS/SOL. Only variants with *P* values < 5×10^−7^ and MAF > 0.5% were included. Previously reported marginal genetic effects, gene-sex interactions, or joint effects within 1Mb flanking regions were shown. Genome coordinates presented were based on GRCh38.

**Table S3.** Significant association regions with LDL cholesterol levels from variant set tests in a 20kb sliding window analysis in HCHS/SOL. Genome coordinates presented were based on GRCh38.

**Table S4.** Significant association regions with LDL cholesterol levels from single-variant meta-analysis combining stochastic summary statistics from HCHS/SOL, ARIC EA and ARIC AA. Only variants with MAF > 0.5% were included. Genome coordinates presented were based on GRCh38.

**Table S5.** Significant association regions with LDL cholesterol levels from variant set meta-analysis in a 20kb sliding window analysis after combining stochastic summary statistics from HCHS/SOL, ARIC EA and ARIC AA. Genome coordinates presented were based on GRCh38.

**Table S6.**
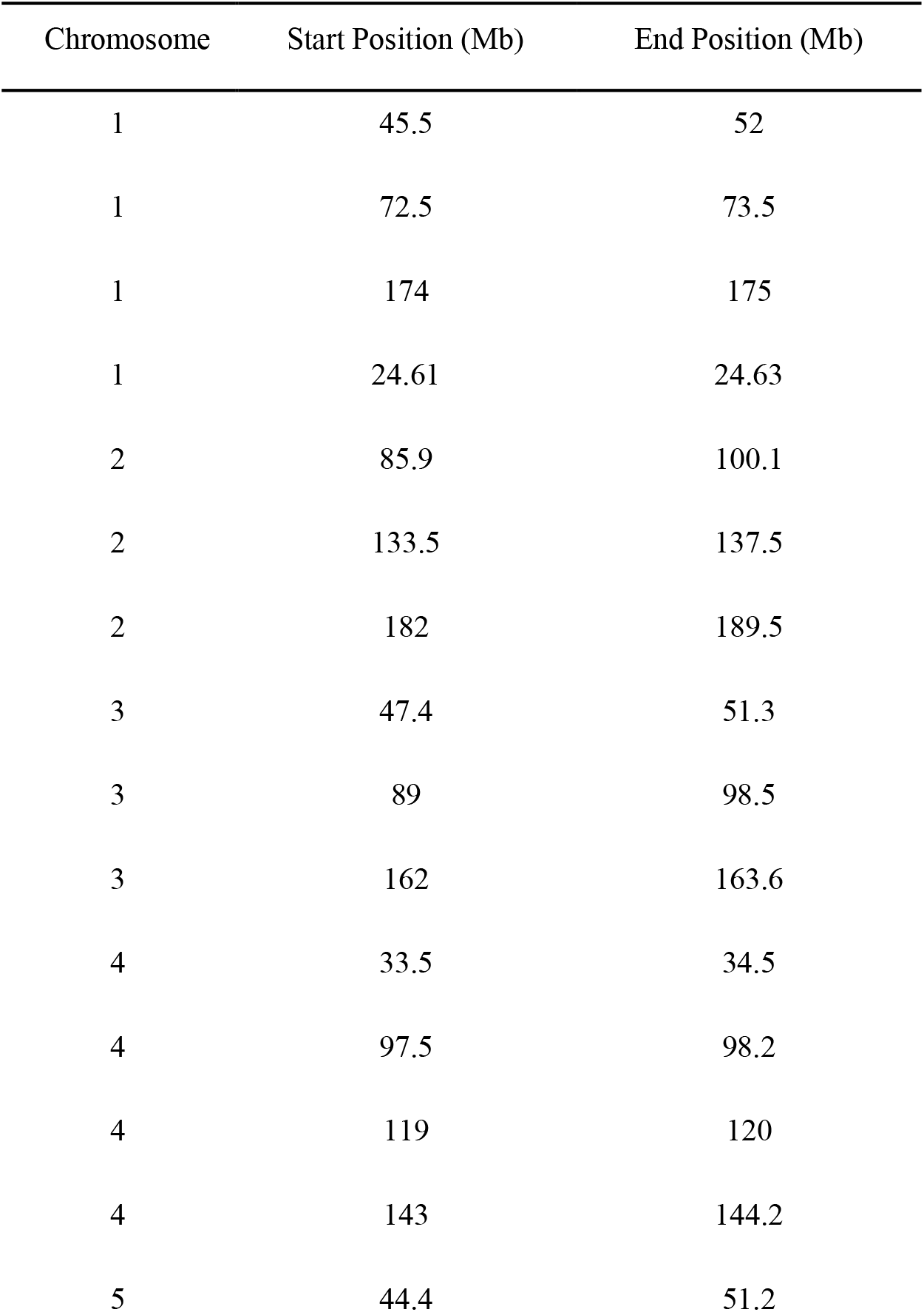
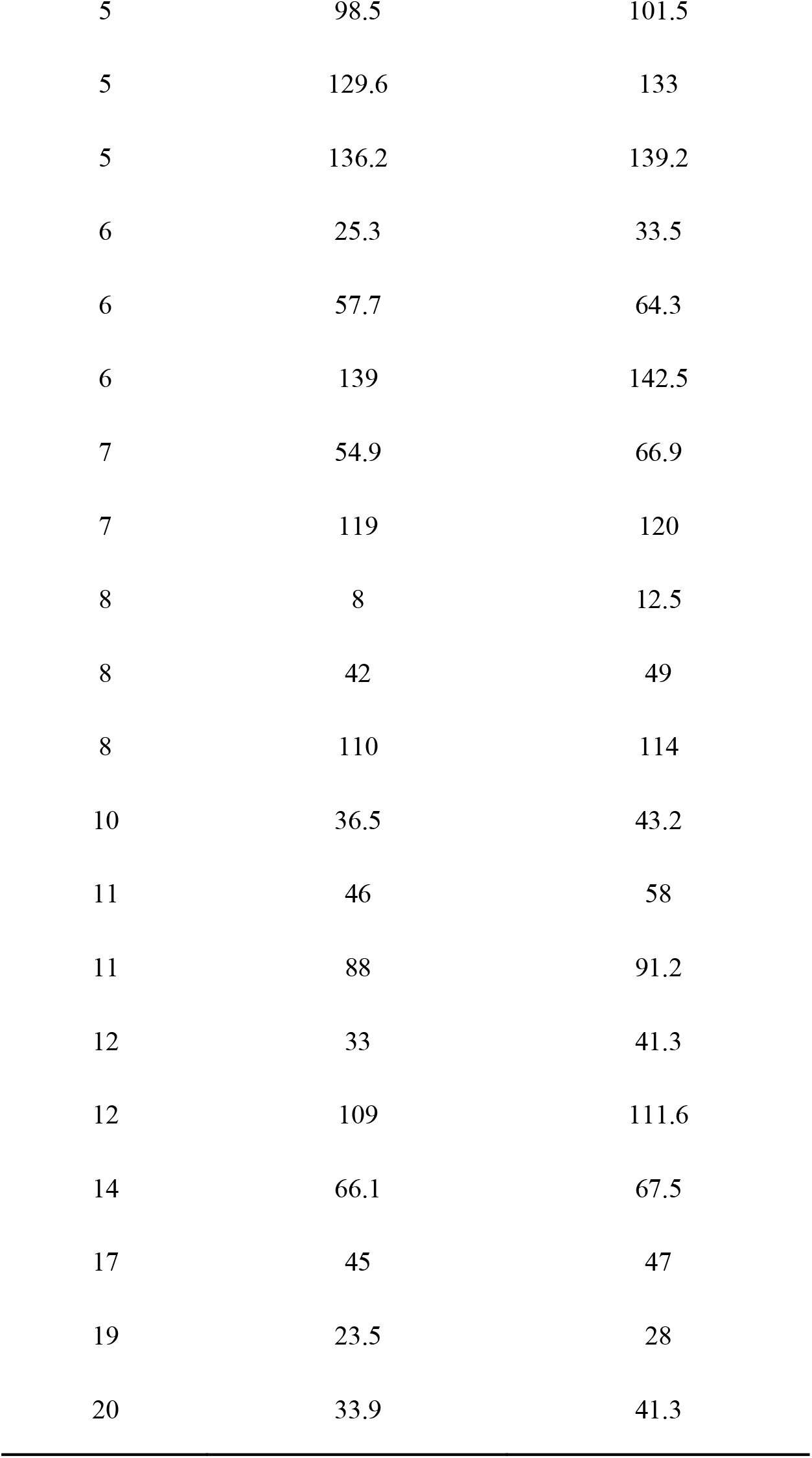
Regions excluded from LD score regression due to long-range LD on the human genome. Genome coordinates presented were based on GRCh38.

| Chromosome | Start Position (Mb) | End Position (Mb) |
| --- | --- | --- |
| 1 | 45.5 | 52 |
| 1 | 72.5 | 73.5 |
| 1 | 174 | 175 |
| 1 | 24.61 | 24.63 |
| 2 | 85.9 | 100.1 |
| 2 | 133.5 | 137.5 |
| 2 | 182 | 189.5 |
| 3 | 47.4 | 51.3 |
| 3 | 89 | 98.5 |
| 3 | 162 | 163.6 |
| 4 | 33.5 | 34.5 |
| 4 | 97.5 | 98.2 |
| 4 | 119 | 120 |
| 4 | 143 | 144.2 |
| 5 | 44.4 | 51.2 |
| 5 | 98.5 | 101.5 |
| 5 | 129.6 | 133 |
| 5 | 136.2 | 139.2 |
| 6 | 25.3 | 33.5 |
| 6 | 57.7 | 64.3 |
| 6 | 139 | 142.5 |
| 7 | 54.9 | 66.9 |
| 7 | 119 | 120 |
| 8 | 8 | 12.5 |
| 8 | 42 | 49 |
| 8 | 110 | 114 |
| 10 | 36.5 | 43.2 |
| 11 | 46 | 58 |
| 11 | 88 | 91.2 |
| 12 | 33 | 41.3 |
| 12 | 109 | 111.6 |
| 14 | 66.1 | 67.5 |
| 17 | 45 | 47 |
| 19 | 23.5 | 28 |
| 20 | 33.9 | 41.3 |

## Supplementary Figures

**Figure S1.**
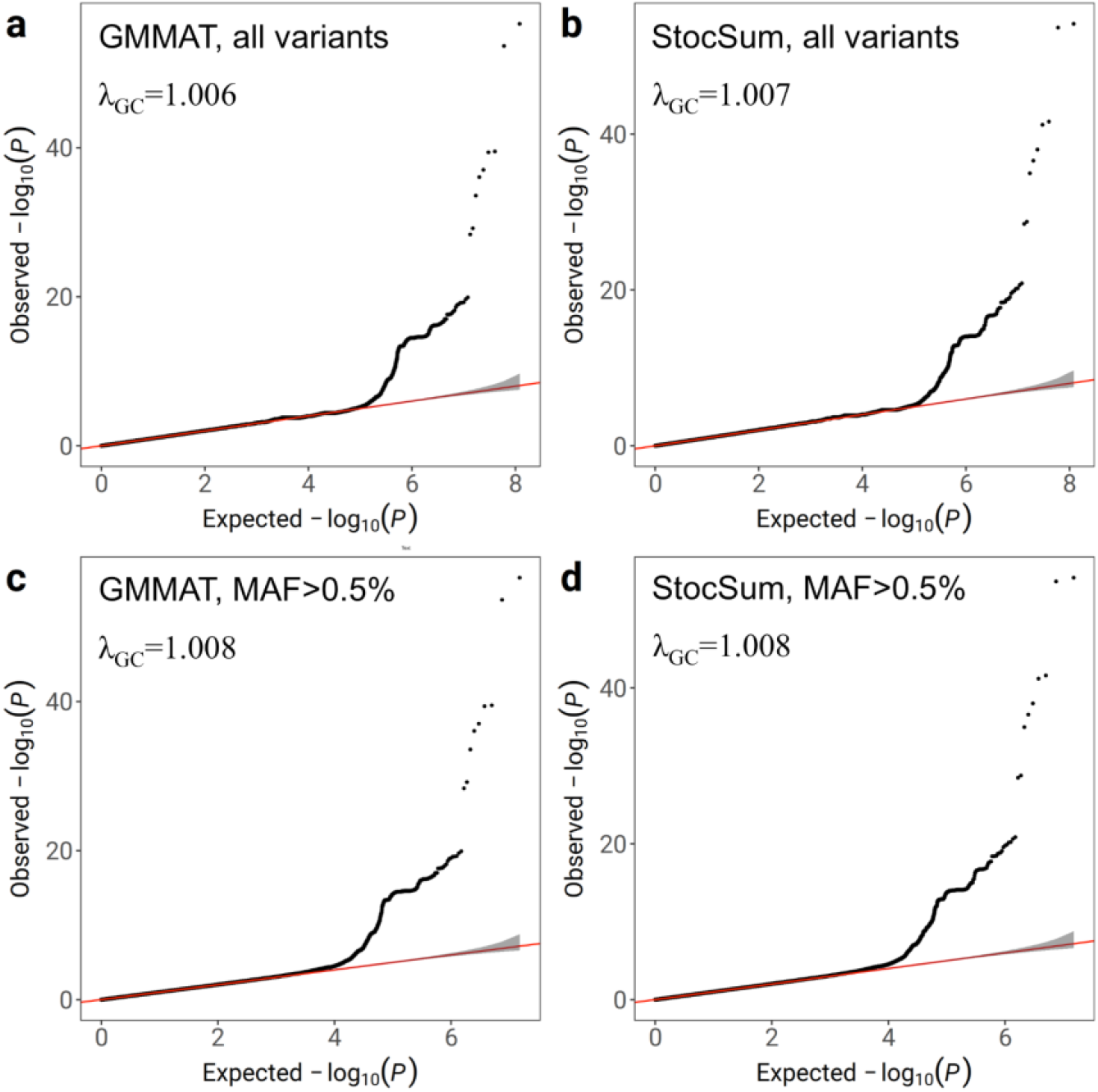
Quantile-quantile (Q-Q) plots of *P* values from single-variant tests on LDL cholesterol levels using GMMAT and StocSum in HCHS/SOL. The number of random vector replicates *B* in StocSum was set to 1,000. a, GMMAT *P* values from all variants. b, StocSum *P* values from all variants. c, GMMAT *P* values from variants with MAF > 0.5%. d, StocSum *P* values from variants with MAF > 0.5%. The gray shaded areas in the Q-Q plots represent 95% confidence intervals under the null hypothesis of no genetic associations.

**Figure S2.**
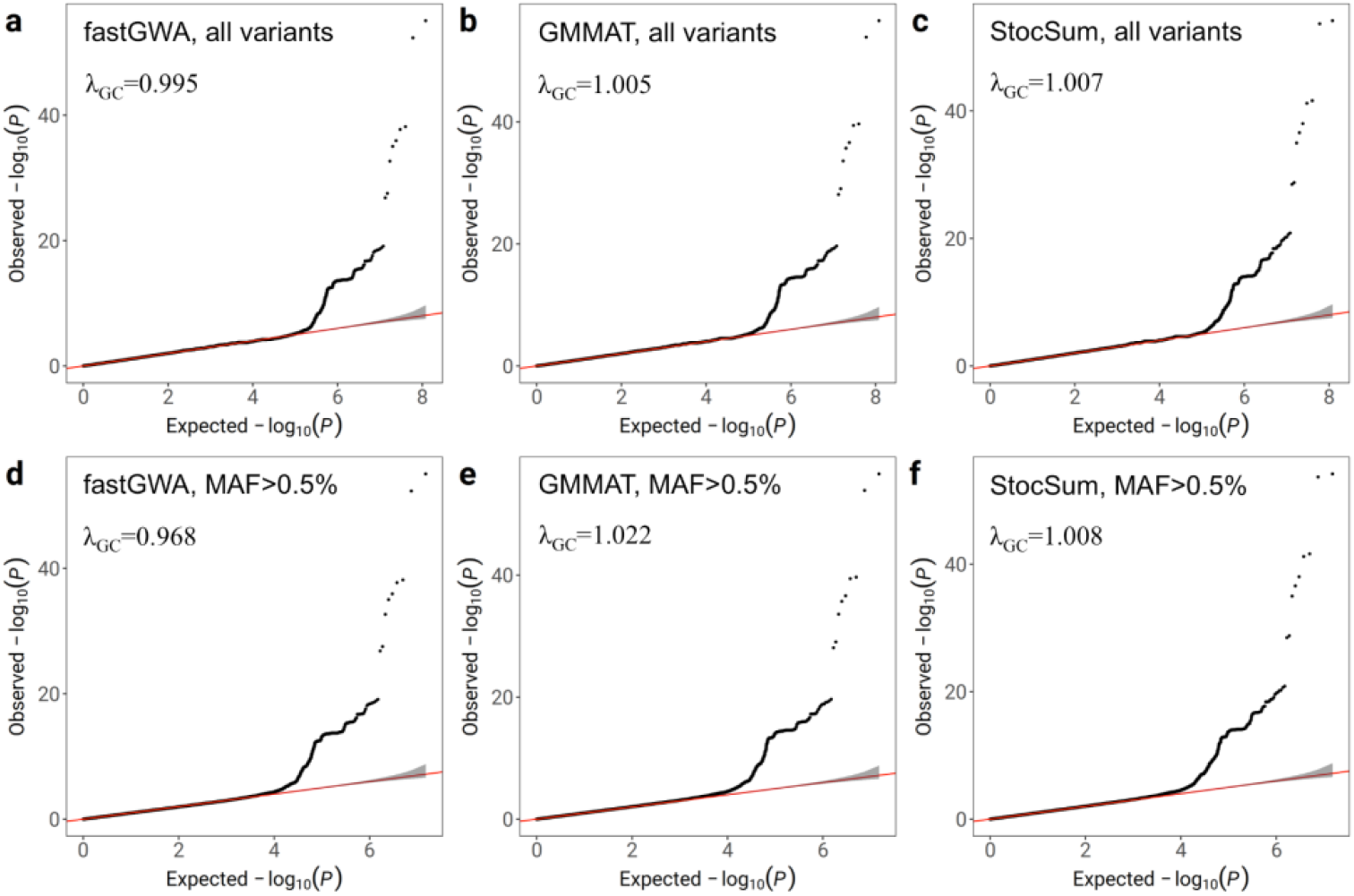
Quantile-quantile (Q-Q) plots of *P* values from single-variant tests on LDL cholesterol levels using fastGWA, GMMAT, and StocSum in HCHS/SOL. The number of random vector replicates *B* in StocSum was set to 1,000. a, fastGWA *P* values from all variants. b, GMMAT *P* values from all variants. c, StocSum *P* values from all variants. d, fastGWA *P* values from variants with MAF > 0.5%. e, GMMAT *P* values from variants with MAF > 0.5%. f, StocSum *P* values from variants with MAF > 0.5%. The gray shaded areas in the Q-Q plots represent 95% confidence intervals under the null hypothesis of no genetic associations.

**Figure S3.**
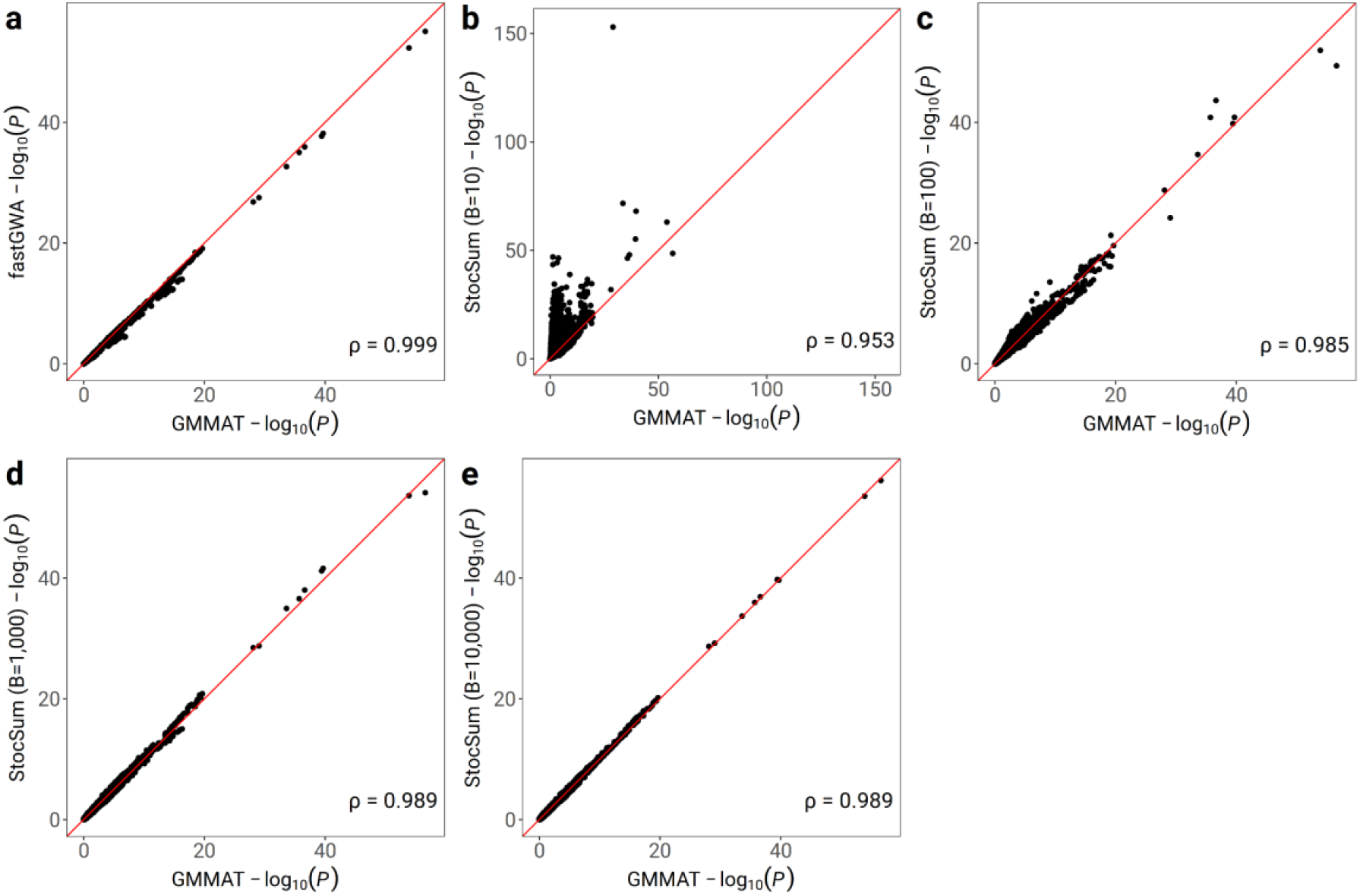
Comparison of *P* values from single-variant tests on LDL cholesterol levels using fastGWA, GMMAT, and StocSum in HCHS/SOL. a, comparison of *P* values from GMMAT and fastGWA. b-e, comparisons of *P* values from GMMAT and StocSum with the number of random vector replicates *B* being equal to 10 (b), 100 (c), 1,000 (d), and 10,000 (e). The red line denotes the reference line of equality. Spearman’s rank correlation coefficients are shown at the bottom right. The data used in this test consisted of 120M variants from 7,297 individuals in HCHS/SOL.

**Figure S4.**
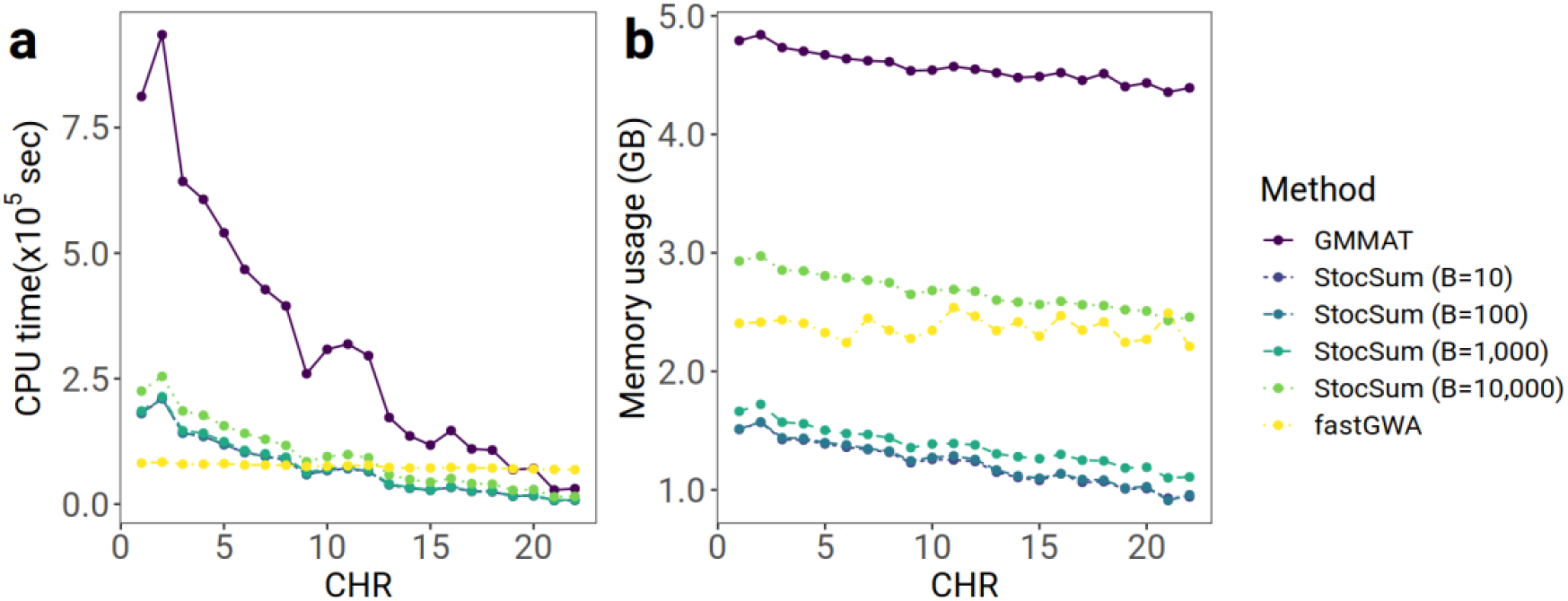
Comparison of CPU time and memory usage from fastGWA, GMMAT and StocSum in single-variant tests. a, CPU time. The x axis represents the chromosome numbers and the y axis represents the CPU time in 10^5^ seconds. For GMMAT, the CPU time consists of fitting the null model and conducting the association test. For StocSum, the CPU time is the sum of four steps: fitting the null model, generating the random vectors, computing the single-variant score statistics and the stochastic summary statistics, and computing the *P* values. b, Memory usage. The x axis represents the chromosome numbers and the y axis represents the memory footprint per thread in GB. The data used in this test consisted of 120M variants from 7,297 individuals in HCHS/SOL. All tests were performed on a high-performance computing server, with 64 threads running in parallel.

**Figure S5.**
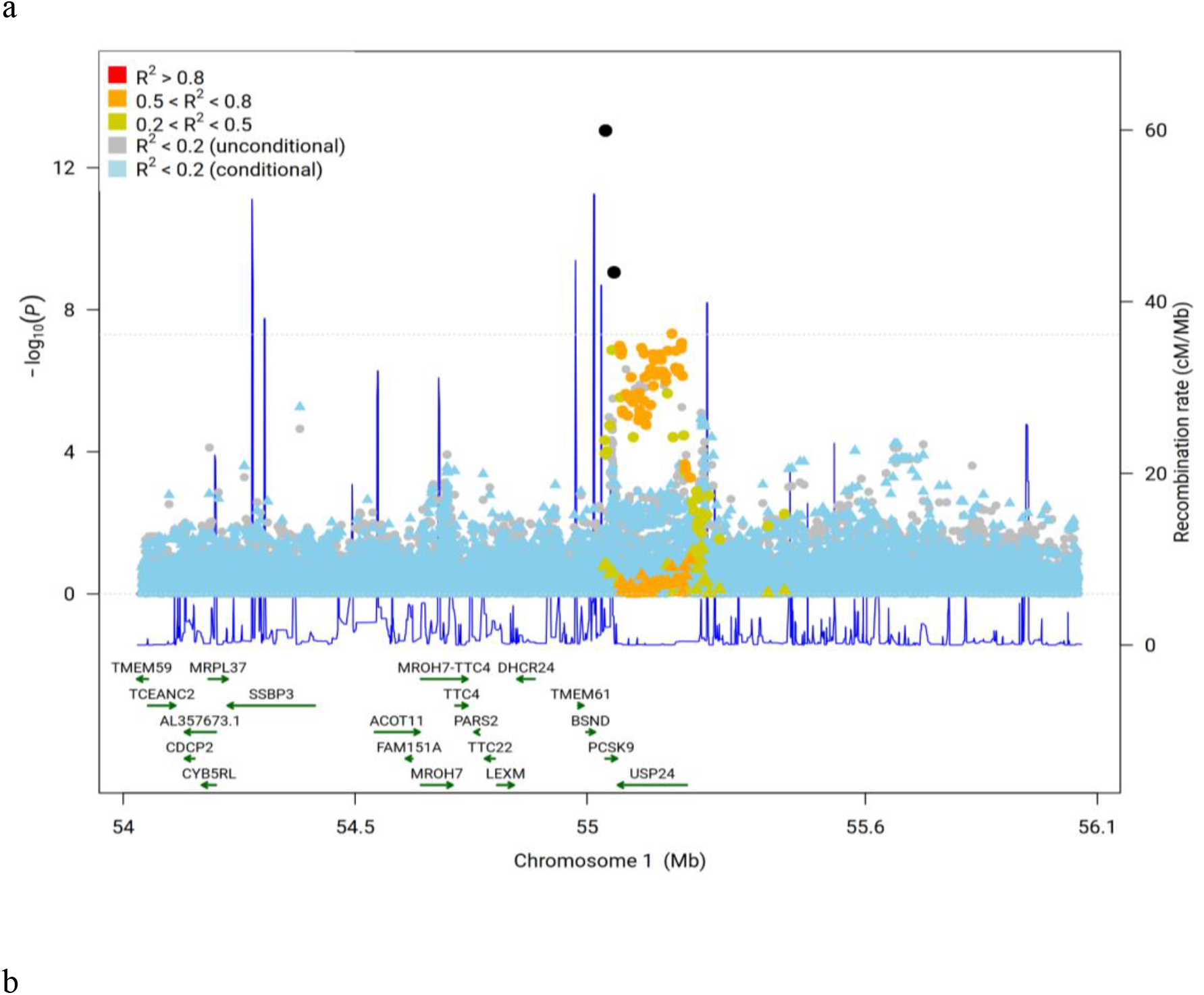
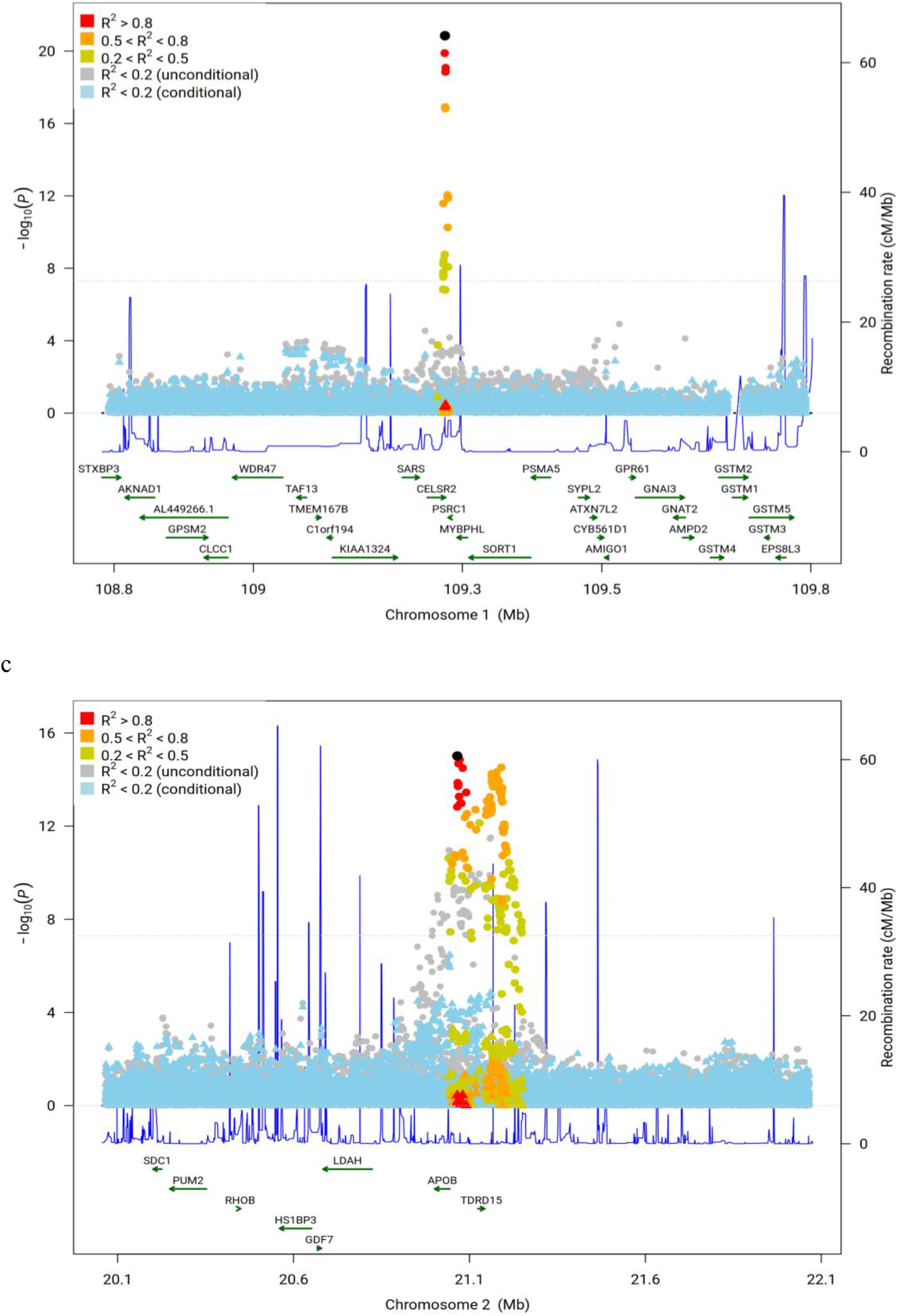
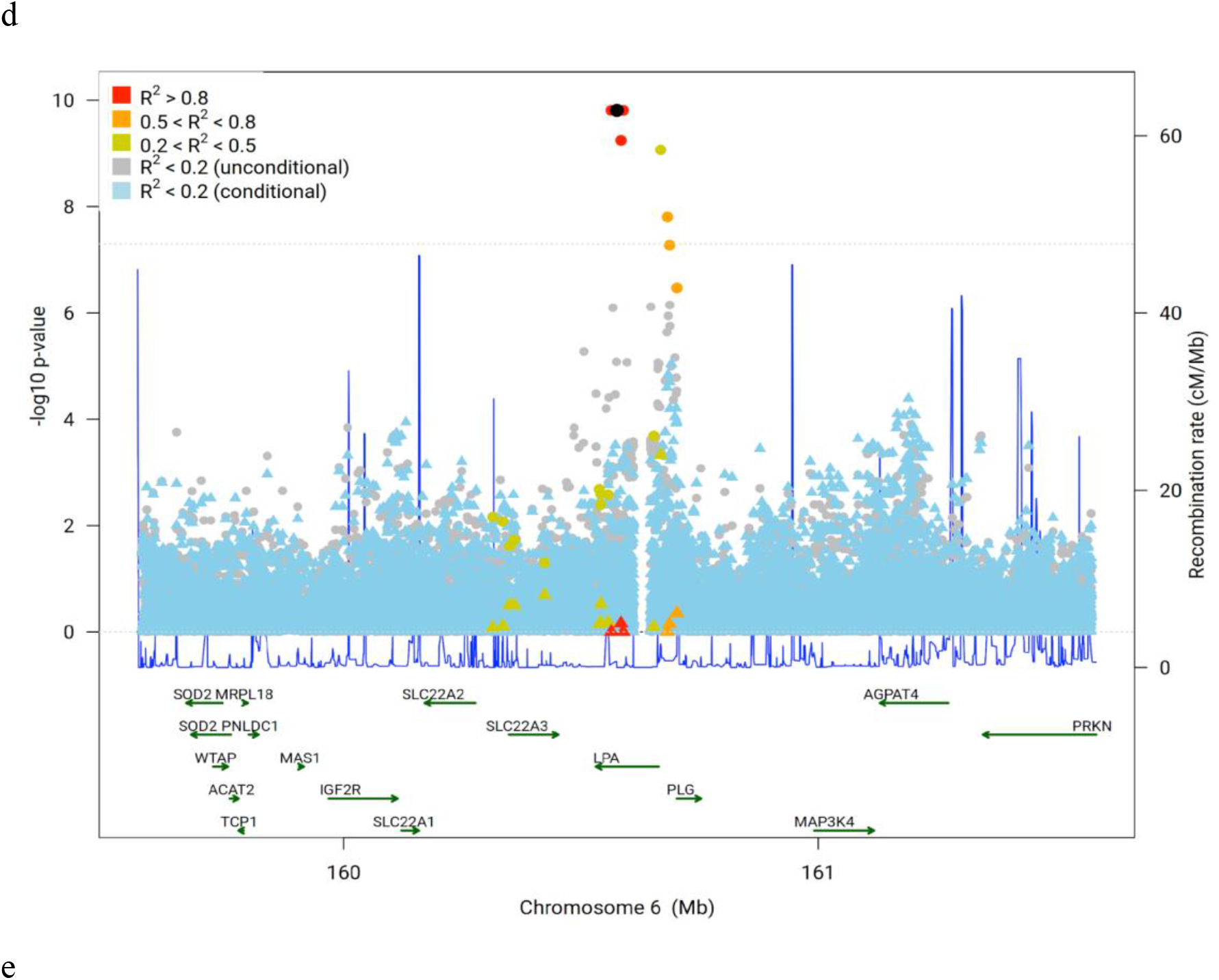
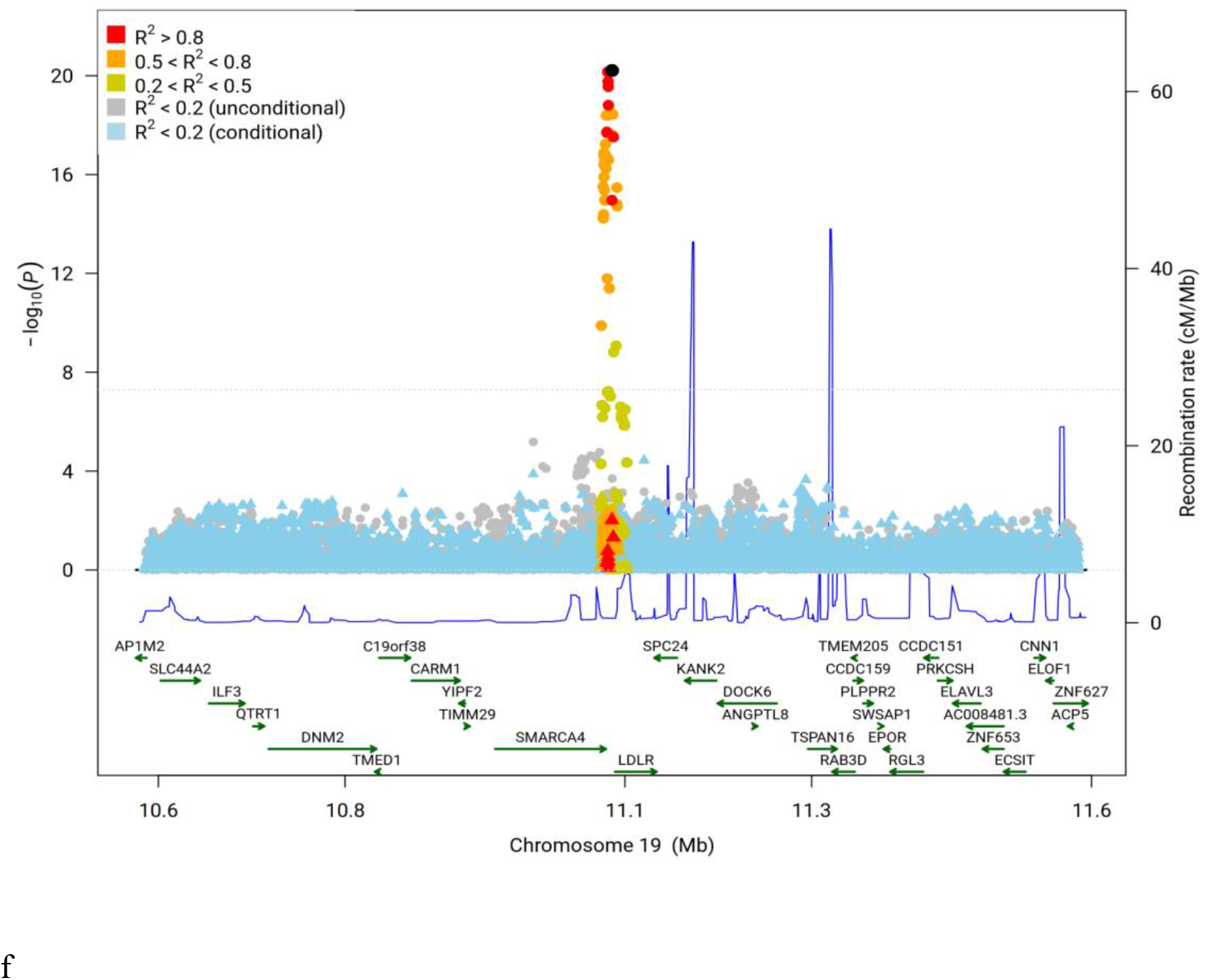
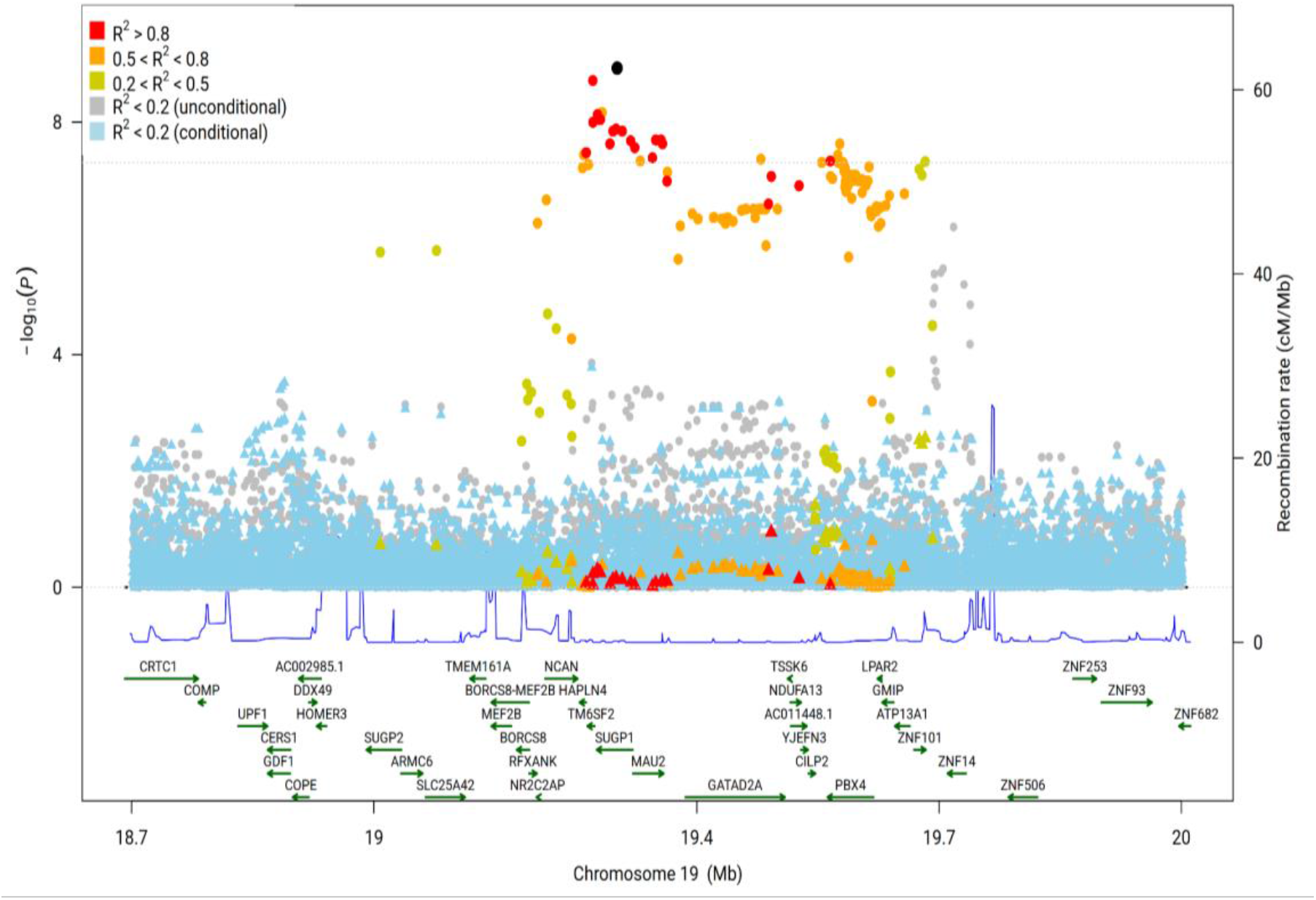
Regional plots of StocSum conditional association test results. a, *PCSK9* gene region with association variants chr1:55039974 (rs28362263) and chr1:55058182 (rs28362263). b, *CELSR2* gene region with the sentinel variant chr1:109274968 (rs562338). c, *APOB* gene region with the sentinel variant chr2:21065449 (rs562338). d, *LPA* gene region with the sentinel variant chr6:160576086 (rs10455872). e, *LDLR* gene region with the sentinel variant chr19:11086210 (rs8106503). f, *SUGP1* gene region with the sentinel variant chr19:19301236 (rs57915152). Association variants are highlighted in black dots. Original single-variant test *P* values are shown in dots and conditional *P* values are shown in triangles. Variants in four LD categories are shown in different colors based on the maximum squared correlation to the sentinel variant and the secondary association variant calculated in HCHS/SOL if there are two association variants (a), or the squared correlation to the sentinel variant in HCHS/SOL if there is only one sentinel association variant (b-f). The horizontal line indicates the genome-wide significance level on the log scale, −log_10_(5 × 10^−8^). The blue curve shows recombination rates from all populations in the 1000 Genome Project.

**Figure S6.**
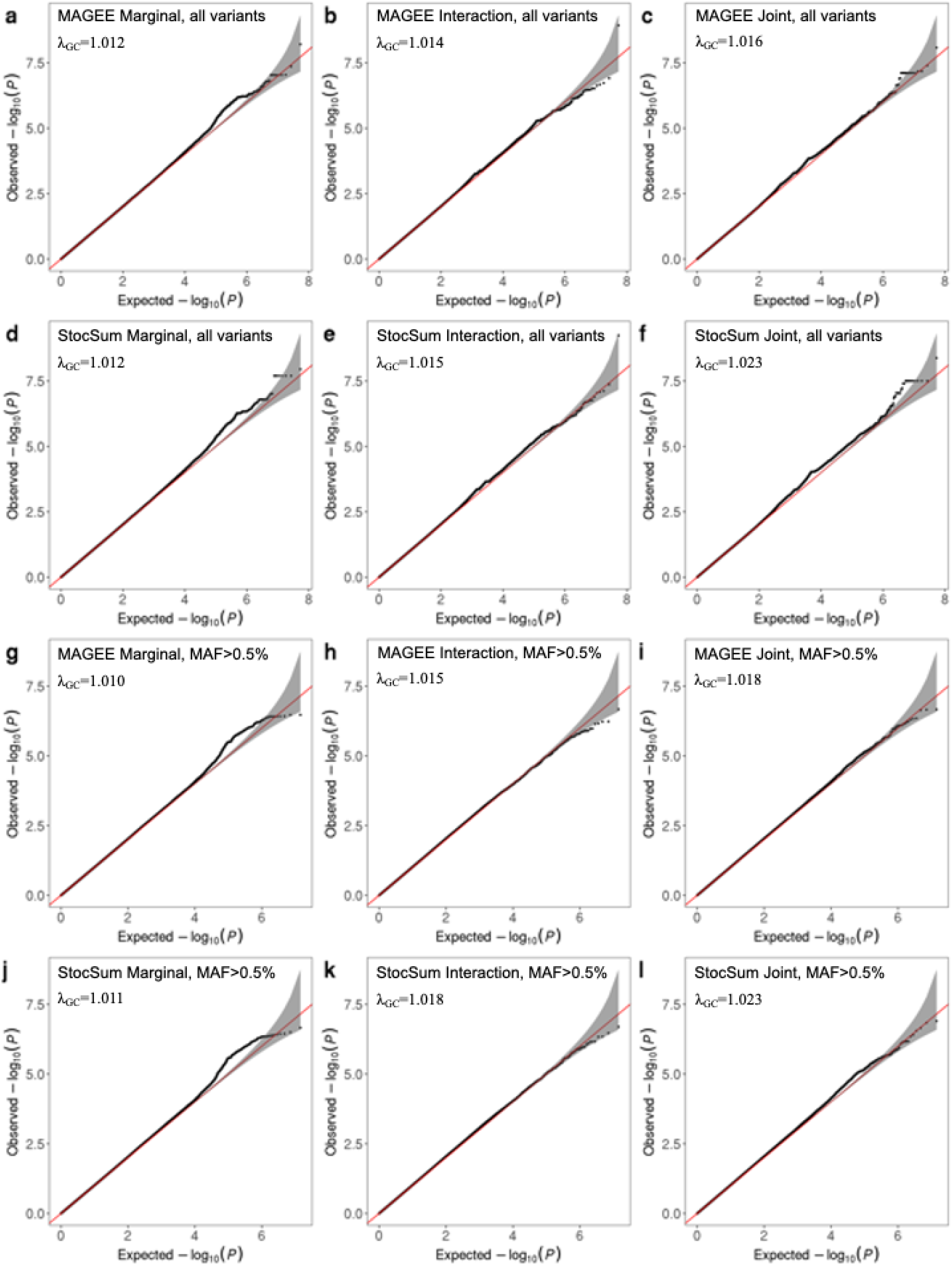
Quantile-quantile (Q-Q) plots of *P* values from gene-sex interaction tests on WHR using MAGEE and StocSum in HCHS/SOL. The number of random vector replicates *B* in StocSum was set to 1,000. a, Marginal *P* values for all variants from MAGEE. b, Interaction *P* values for all variants from MAGEE. c, Joint *P* values for all variants from MAGEE. d, Marginal *P* values for all variants from StocSum. e, Interaction *P* values for all variants from StocSum. f, Joint *P* values for all variants from StocSum. g, Marginal *P* values for variants with MAF > 0.5% from MAGEE. h, Interaction *P* values for variants with MAF > 0.5% from MAGEE. i, Joint *P* values for variants with MAF > 0.5% from MAGEE. j, Marginal *P* values for variants with MAF > 0.5% from StocSum. k, Interaction *P* values for variants with MAF > 0.5% from StocSum. l, Joint *P* values for variants with MAF > 0.5% from StocSum. The gray shaded areas in the Q-Q plots represent 95% confidence intervals under the null hypothesis of no genetic associations and/or gene-sex interactions.

**Figure S7.**
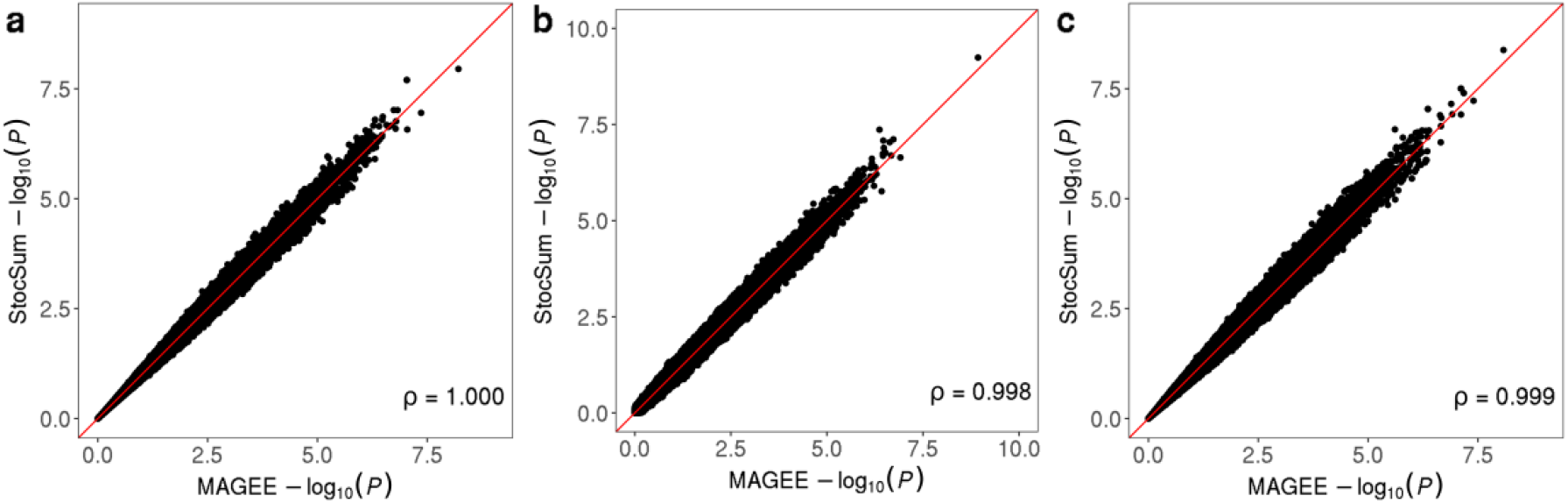
Comparison of *P* values from single-variant gene-sex interaction tests on WHR using MAGEE and StocSum in HCHS/SOL. a, comparison of marginal genetic effect test *P* values. b, comparison of gene-sex interaction test *P* values. c, comparison of joint test *P* values. The x axis and the y axis represent −log_10_(*P*) using MAGEE and StocSum, respectively. The red line denotes the reference line of equality. Spearman’s rank correlation coefficients are shown at the bottom right.

**Figure S8.**
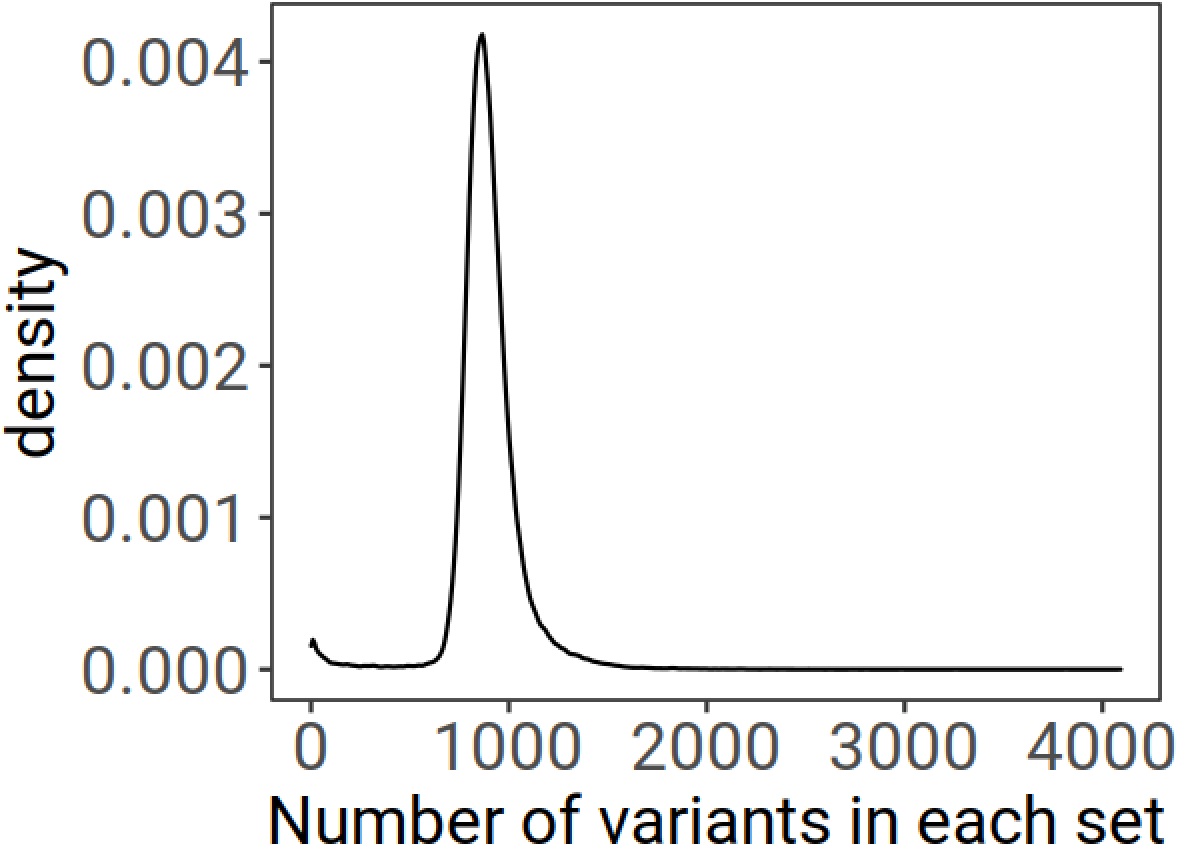
A density plot showing the distribution of variant numbers in each set in a 20 kb sliding window analysis on LDL cholesterol levels in HCHS/SOL.

**Figure S9.**
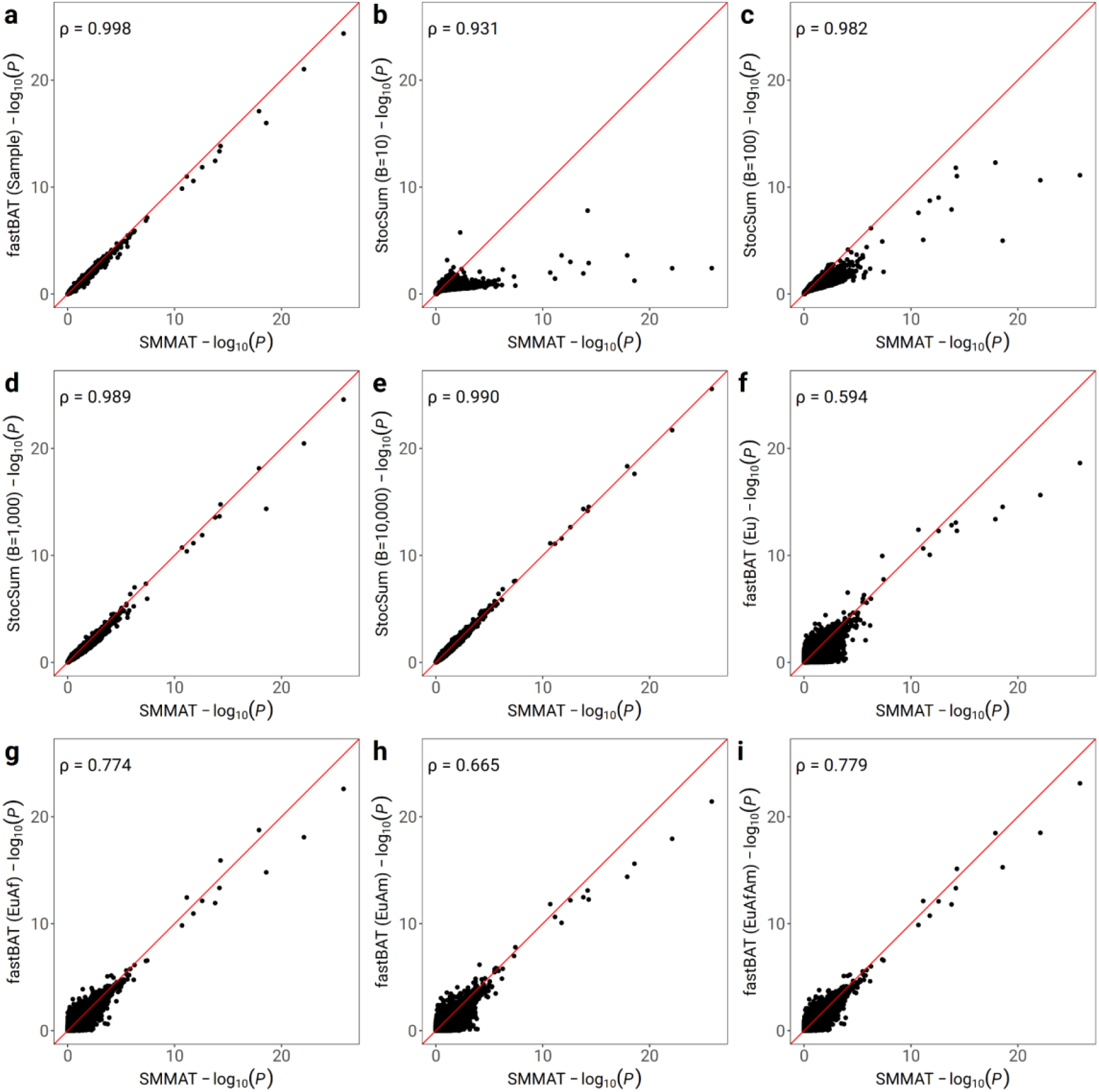
Comparison of *P* values from variant set tests in a 20 kb sliding window analysis on LDL cholesterol levels using fastBAT, SMMAT, and StocSum in HCHS/SOL. The x axis represents the −log_10_(*P*) from variant set tests using SMMAT on individual-level data and the y axis represents the −log_10_(*P*) from variant set tests using StocSum or fastBAT. a, fastBAT with an internal reference panel using the HCHS/SOL study samples (fastBAT (Sample)). b-e, StocSum with the number of random vector replicates *B* being equal to 10 (b), 100 (c), 1,000 (d) and 10,000 (e). f-i, fastBAT with external reference panels from 1000 Genomes using European (fastBAT (Eu)) (f), European and African (fastBAT (EuAf)) (g), European and American (fastBAT (EuAm)) (h), and European, African, and American (fastBAT (EuAfAm)) (i) populations. The red line denotes the reference line of equality. The data used in this test consisted of 120M variants from 7,297 individuals in HCHS/SOL. Spearman’s rank correlation coefficients are shown at the top left.

**Figure S10.**
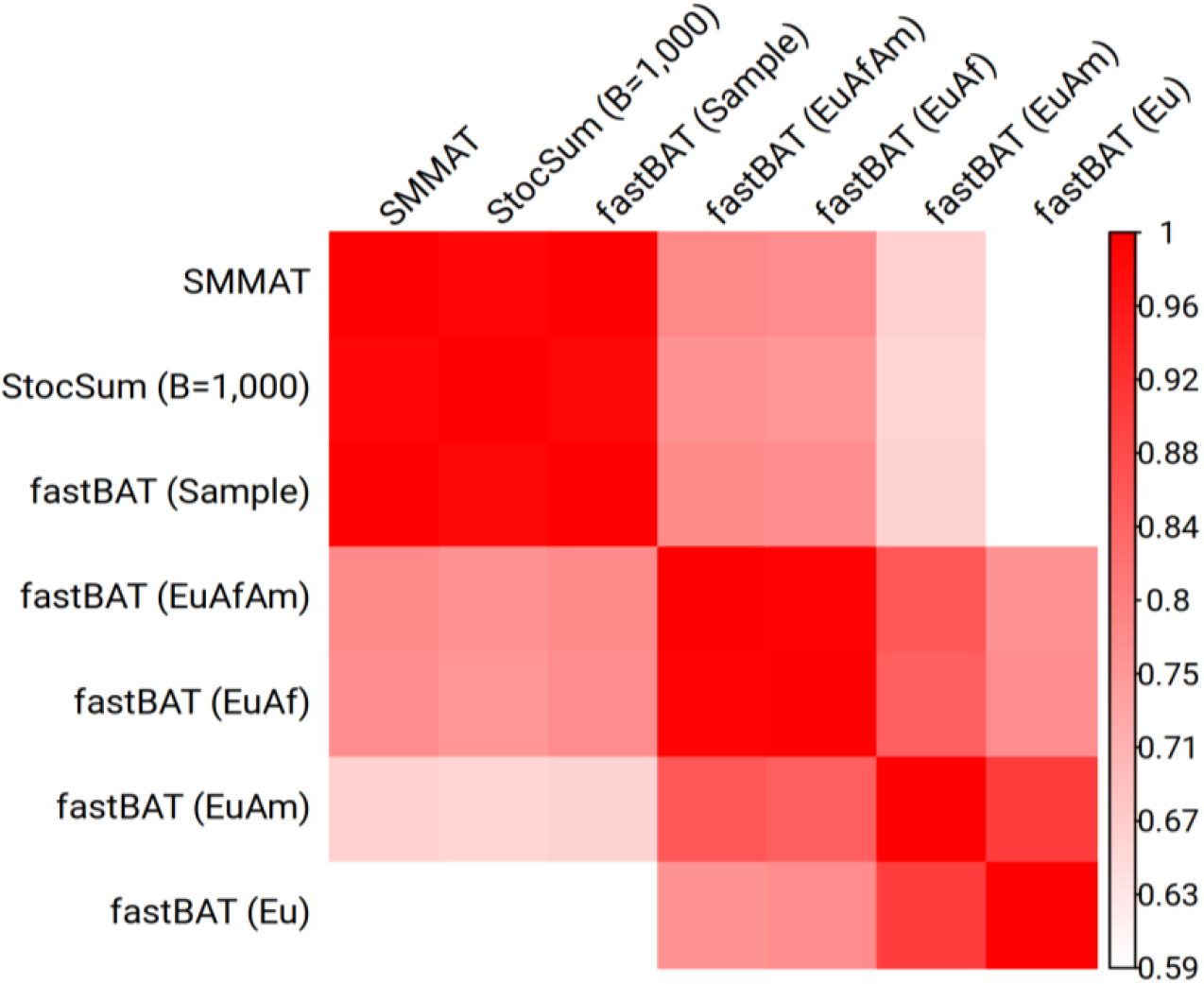
Heatmap showing Spearman’s rank correlation coefficients of *P* values from variant set tests in a 20 kb sliding window analysis on LDL cholesterol levels using fastBAT, SMMAT, and StocSum in HCHS/SOL. For fastBAT, we used an internal reference panel using the HCHS/SOL study samples (fastBAT (Sample)), as well as four external reference panels from 1000 Genomes (Eu, EuAf, EuAm, EuAfAm).

**Figure S11.**
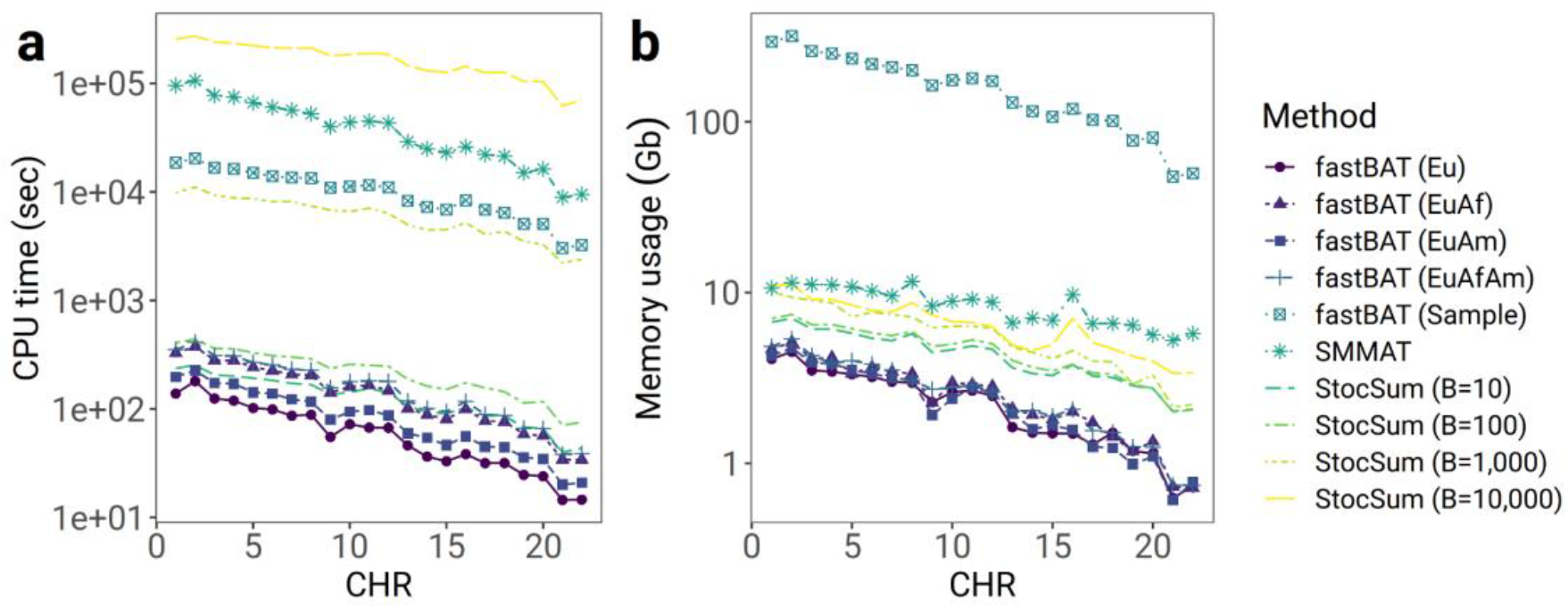
Comparison of CPU time and memory usage from fastBAT, SMMAT, and StocSum in variant set tests in a 20 kb sliding window analysis on LDL cholesterol levels in HCHS/SOL. a, CPU time. The x axis represents the chromosome numbers and the y axis represents the CPU time on the logarithmic scale. The CPU time only includes the step of computing the *P* values, assuming corresponding summary statistics have been computed in single-variant tests. b, Memory usage. The x axis represents the chromosome numbers and the y axis represents the memory footprint per thread in GB on the logarithmic scale. The data used in this test consisted of 120M variants from 7,297 individuals in HCHS/SOL. All tests were performed on a high-performance computing server, with a single thread for each chromosome.

**Figure S12.**
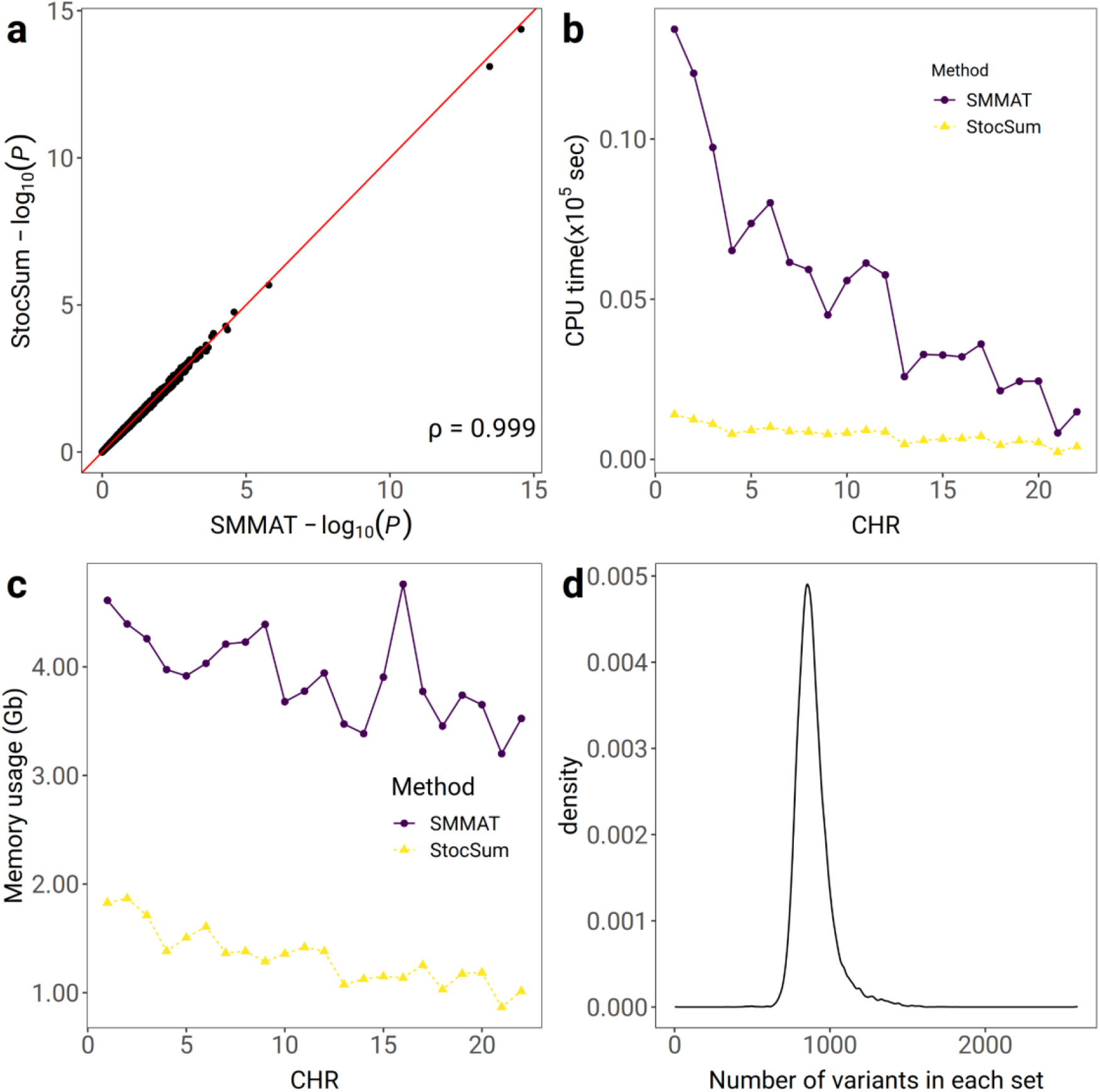
Comparison of SMMAT and StocSum variant set tests in a non-sliding-window analysis on LDL cholesterol levels in HCHS/SOL. The variant sets were defined by merging chromatin loops of H3K27ac HiChIP interaction in the GM12878 cell line. There are a total of 17,224 paired regions, each as a variant set, including two 10kb windows which may not be located in close proximity on the primary structure of DNA and not typically covered using fixed-size sliding windows. a, comparison of *P* values from SMMAT and StocSum with the number of random vector replicates *B* being equal to 1,000. The x axis and the y axis represent the −log_10_(*P*) from variant set tests using SMMAT and StocSum, respectively. The red line denotes the reference line of equality. b, comparison of CPU time between SMMAT and StocSum. The x axis represents the chromosome numbers and the y axis represents the CPU time in 10^5^ seconds. For SMMAT and StocSum, the CPU time only includes the step of computing the *P* values, assuming corresponding summary statistics have been computed in single-variant tests. c, comparison of memory usage between SMMAT and StocSum. The x axis represents the chromosome numbers and the y axis the memory footprint per thread in GB. d, a density plot showing the distribution of variant numbers in each set.

**Figure S13.**
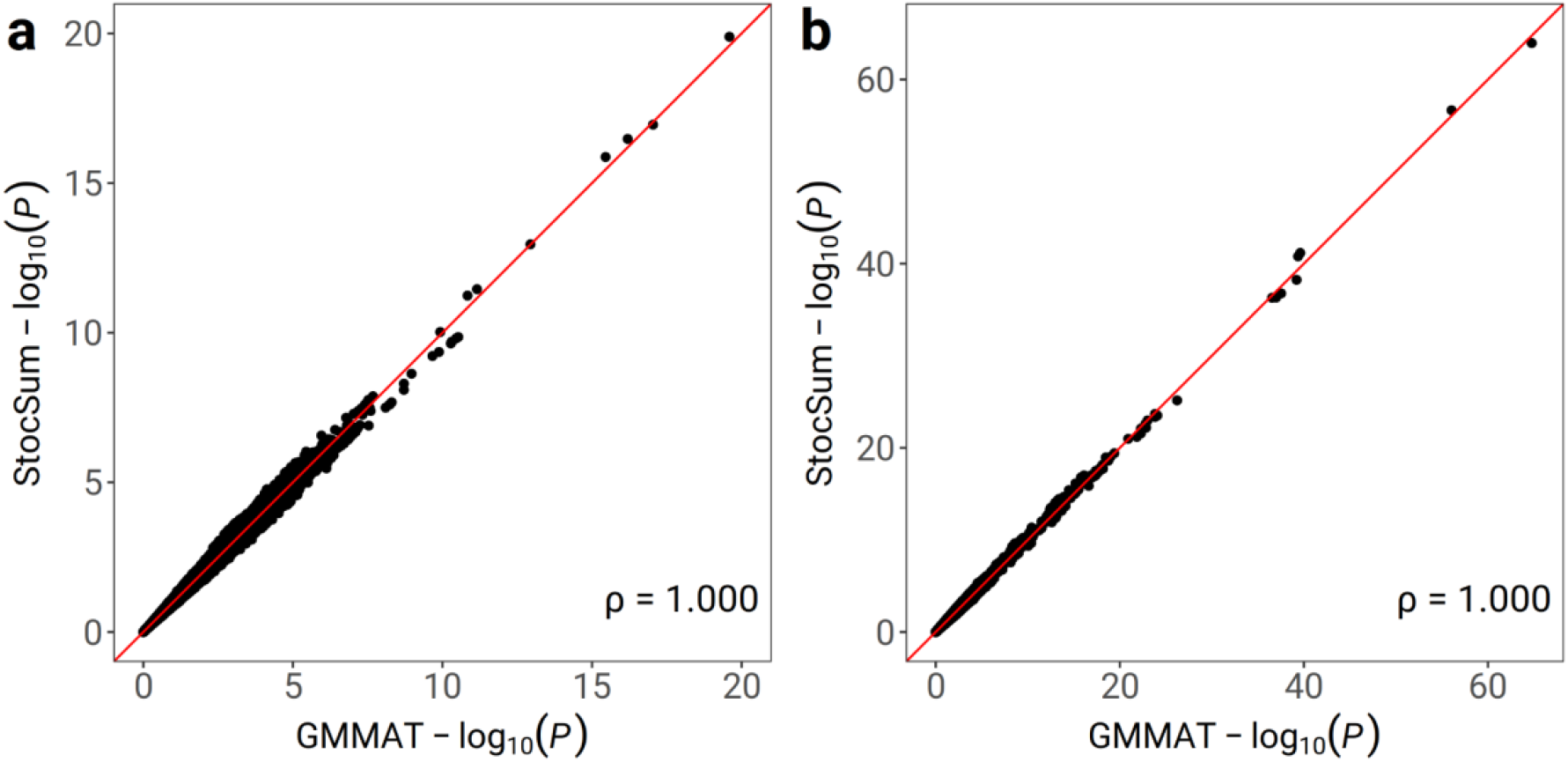
Comparison of *P* values from single-variant tests on longitudinal LDL cholesterol levels using GMMAT and StocSum in ARIC AA (a) and ARIC EA (b). The ARIC AA data used in this test consisted of 70M variants and 7,514 observations from 2,045 individuals. The ARIC EA data used in this test consisted of 92M variants and 26,668 observations from 6,327 individuals. The x axis and the y axis represent the −log_10_(*P*) from single-variant tests using GMMAT and StocSum with the number of random vector replicates *B* being equal to 1,000. The red line denotes the reference line of equality. Spearman’s rank correlation coefficients are shown at the bottom right.

**Figure S14.**
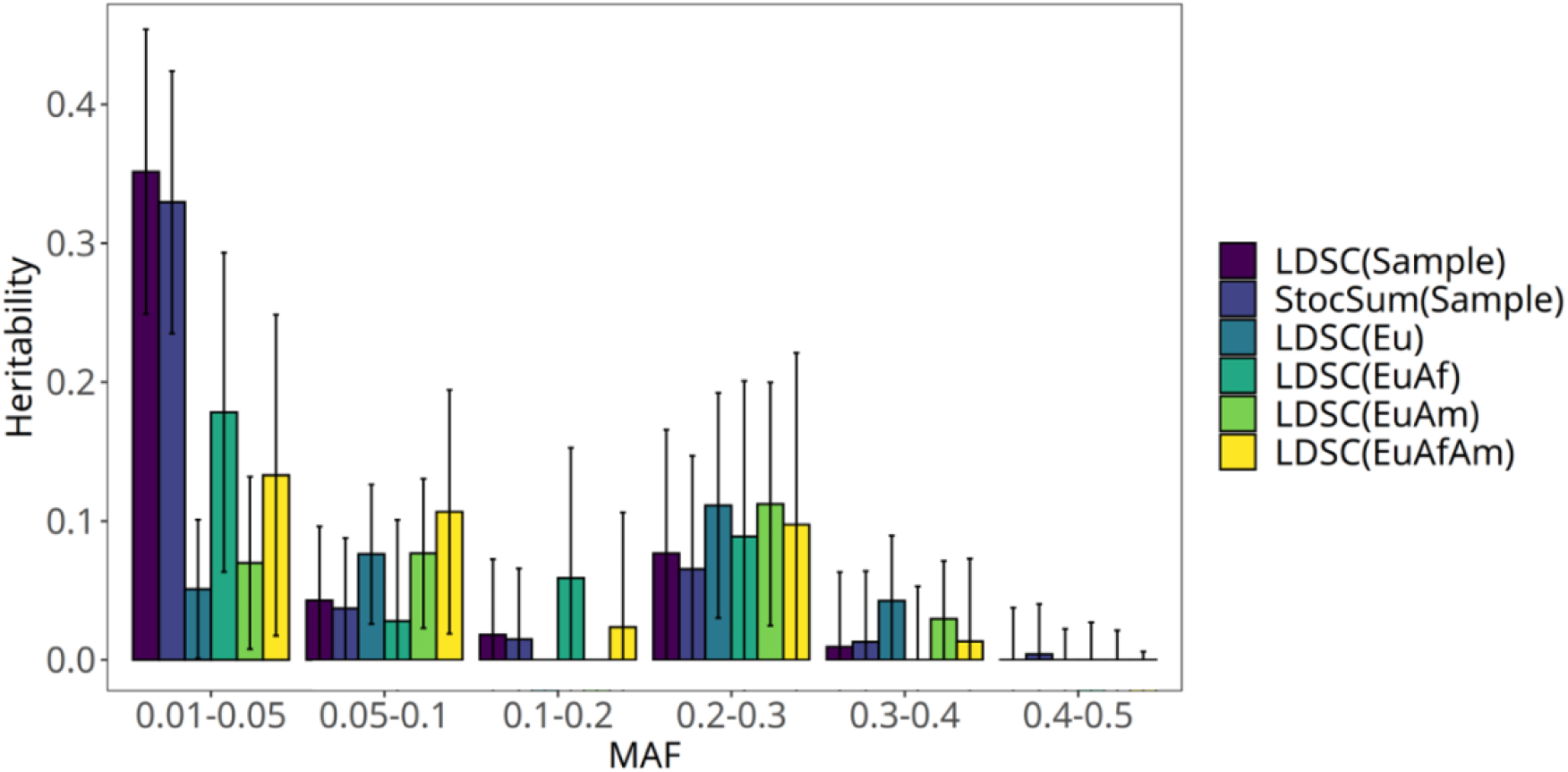
LDL heritability estimates by stratified LDSC and StocSum for different MAF bins. The error bars show point estimates ± standard errors. Negative heritability estimates reported from stratified LDSC were truncated at 0. LD scores for different MAF bins were estimated from LDSC (Sample) and StocSum (Sample) using HCHS/SOL study samples, or LDSC on external reference panels using European, African and/or American populations from the 1000 Genomes Project: LDSC (Eu), LDSC (EuAf), LDSC (EuAm), and LDSC (EuAfAm).

## Notes

### Competing Interest Statement

The authors have declared no competing interest.

### Summary of Updates

An author name updated

